# ARF6 as a novel activator of HIF-2α in pulmonary arterial hypertension

**DOI:** 10.1101/2023.09.15.557917

**Authors:** Adam L. Fellows, Chien-Nien Chen, Chongyang Xie, Nayana Iyer, Lukas Schmidt, Xiaoke Yin, Manuel Mayr, Andrew Cowburn, Lan Zhao, Beata Wojciak-Stothard

**Author notes:** Correspondence Dr Beata Wojciak-Stothard, National Heart and Lung Institute (NHLI), ICTEM Building, Hammersmith Campus, Imperial College London, Du Cane Road, London W12 0NN, or Dr Adam Fellows, National Heart and Lung Institute (NHLI), ICTEM Building, Hammersmith Campus, Imperial College London, Du Cane Road, London W12 0NN.

## Abstract

ADP-ribosylation factor 6 (ARF6), a GTPase associated with cancer metastasis, is activated in the lung endothelium in pulmonary arterial hypertension (PAH). To identify ARF6-regulated pathways relevant to PAH, we performed a state-of-the-art proteomic analysis of human pulmonary artery endothelial cells (HPAECs) overexpressing the wildtype, constitutively active, fast-cycling and dominant negative mutants of ARF6. The analysis revealed a novel link of ARF6 with hypoxia-inducible factor (HIF), in addition to endocytotic vesicle trafficking, cell proliferation, angiogenesis, oxidative stress and lipid metabolism. Active ARF6 markedly increased expression and activity of HIF-2, critical in PAH, with HIF-1 relatively unaffected. Hypoxic ARF6 activation was a prerequisite for HIF-2 activation and HIF-dependent gene expression in HPAECs, PAH blood-derived late outgrowth endothelial colony forming cells (ECFCs) and hypoxic mouse lungs *in vivo*. A novel ARF6 inhibitor, chlortetracycline (CTC), reduced hypoxia-induced HIF-2 activation, proliferation and angiogenesis in HPAECs and reduced HIF-2 expression in lung and heart tissues of hypoxic mice. PAH ECFCs showed elevated expression and activity of ARF6 and HIF2, which was attenuated by CTC. Oral CTC attenuated development of PH in chronically hypoxic mice. In conclusion, we are first to demonstrate a key role of ARF6 in the regulation of HIF-2α activation *in vitro* and *in vivo* and show that HIF-2α, a master-regulator of vascular remodelling in PAH, can be targeted by a clinically approved antibiotic chlortetracycline.

## INTRODUCTION

Pulmonary arterial hypertension (PAH) is a rare disease characterised by gradual narrowing of small and medium pulmonary arteries, ultimately leading to right heart failure and premature death (1). Pulmonary endothelial dysfunction, manifested by increased proliferation, resistance to apoptosis, angiogenesis and endothelial-to-mesenchymal transition, largely driven by hypoxia-inducible factor-2 (HIF-2), plays a critical role in initiation and progression of PAH (2, 3).

ADP-ribosylation factor 6 (ARF6) has been implicated in pulmonary vascular remodelling in PAH (4) but signalling mechanisms involved are not fully understood.

ARF6 is a member of ARF family of six proteins (ARF1-6), belonging the Ras superfamily of small GTPases (5). Like other ARFs, ARF6 oscillates between an “active” GTP-bound state and an “inactive” GDP-bound state, with GTP binding and hydrolysis mediated by guanine nucleotide exchange factors (GEFs) and GTPase-activating proteins (GAPs), respectively. Whilst the GTP-bound, membrane-associated form carries out most of the activities attributed to ARF6, its GDP-bound form together with regulatory proteins may also initiate signalling events (6). ARF6 is primarily known as a regulator of membrane receptor trafficking, vesicle transport, actin remodelling and cell motility in various cell types, including metastatic cancer cells (7). Recent studies suggest that ARF6 may act as a modulator of hypoxic signalling in certain cell types (8) but specific mechanisms have not been delineated.

PAH has a complex and multi-faceted pathophysiology, including perturbed endothelial function and a heightened response to hypoxia. In this study, we aimed to identify key processes affected by ARF6 in human pulmonary endothelial cells, relevant to PAH. We report a novel pathway whereby ARF6 mediates the activation of hypoxia-inducible factor 2α (HIF-2α) in response to low oxygen tension both *in vitro* and *in vivo* and demonstrate effectiveness of a new ARF6 inhibitor, the antibiotic chlortetracycline, in the inhibition of HIF-2 signalling in healthy and PAH endothelial cells and chronic hypoxia mouse model of PAH.

## RESULTS

### Expression and intracellular localization of ARF6 activity mutants in HPAECs

To better understand the effects of ARF6, HPAECs were infected with recombinant adenoviruses to overexpress the wildtype (WT-ARF6), constitutively active ARF6^Q67L^ (CA-ARF6)(9), fast cycling ARF6^T157A^ (FC-ARF6) (10) and dominant negative ARF6^T27N^ (DN-ARF6)(9) (**Fig. 1A**). In CA-ARF6, the GTP binding is resistant to hydrolysis, hence the mutant remains permanently locked into an active, GTP-bound conformation. The FC-ARF6 mutant exhibits spontaneously higher rates of GTP exchange, whilst retaining a capacity for GTP hydrolysis, whilst the DN-ARF6 mutant displays defective GTP binding and is locked into an inactive, GDP-bound state (**Fig. 1B**). Mutant proteins were effectively overexpressed in >90% of infected cells (**Supplementary Figure S1**).

**Figure 1.**
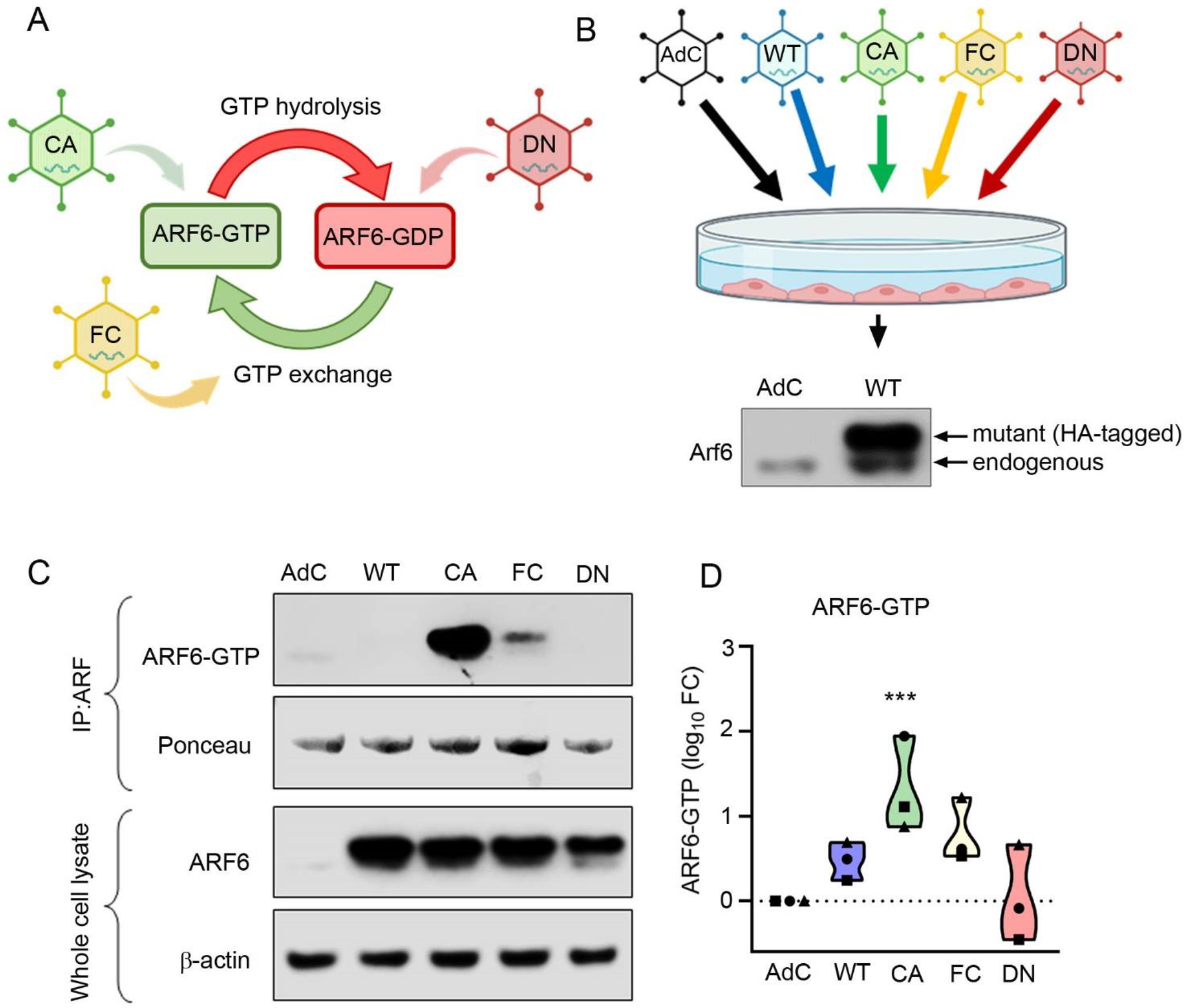
Adenoviral overexpression of ARF6 activity mutants in HPAECs. (**A**) Schematic illustrating effect of ARF6 mutations on GTP hydrolysis and GDP exchange. (**B**) Expression of endogenous and HA-tagged mutant ARF6 (WT, CA, DN and FC) was studied by western blotting in HPAECs 24 hour post-infection. (**C**) Representative western blots and (**D**) a corresponding graph showing changes in the levels of active (GTP-bound) ARF6, total ARF6 and β-actin, as indicated. (**D**) shows fold-changes normalised to the empty adenoviral control (AdC) (n=3 different biological replicates), ***P<0.001; one-way ANOVA with Dunnett’s multiple comparisons test.

To enable a comparative analysis, CA-ARF6 overexpression levels were adjusted to induce ∼10-fold increase in ARF6 activity, corresponding to changes seen in PAH endothelial cells (4), and expression of other ARF6 mutants was equalised to that of CA-ARF6 (**Fig. 1C, D**). Cell fractionation showed that CA-ARF6 localised predominantly to the cell membrane, whilst DN mutant, as predicted, showed mainly cytoplasmic localization (**Supplementary Fig. S2**). WT-ARF6 and FC-ARF6 mutants showed equal distribution between the cell membrane and cytoplasm and were largely absent from cytoskeletal or nuclear fractions (**Supplementary Fig. S2**). ARF6 activity (GTP binding) was highest in CA-ARF6-overexpressing HPAECs, while FC-ARF6 had a more modest effect (**Fig. 1C, D**).

### Proteomic analysis of HPAECs overexpressing ARF6 activity mutants

An unbiased proteomic analysis identified key processes affected by ARF6 activity changes in HPAECs. HPAECs from four different donors overexpressing ARF6 mutants were subjected to a state-of-the-art proteomics workflow including tandem mass tag (TMT) peptide labelling combined with liquid chromatography with tandem mass spectrometry (LC-MS/MS) to ensure high precision coverage and quantitation (11) (Fig. 2A). Robust and equal overexpression of ARF6 protein 24 hours post-adenoviral transduction was confirmed by Western blotting (**Fig. 2B, C**). The number of differentially expressed proteins (DEPs) varied amongst the ARF6 mutants, with WT-ARF6, CA-ARF6, FC-ARF6 and DN-ARF6 mutants generating 3, 14, 19 and 43 DEPs, respectively (**Supplementary Figure S3**).

**Figure 2.**
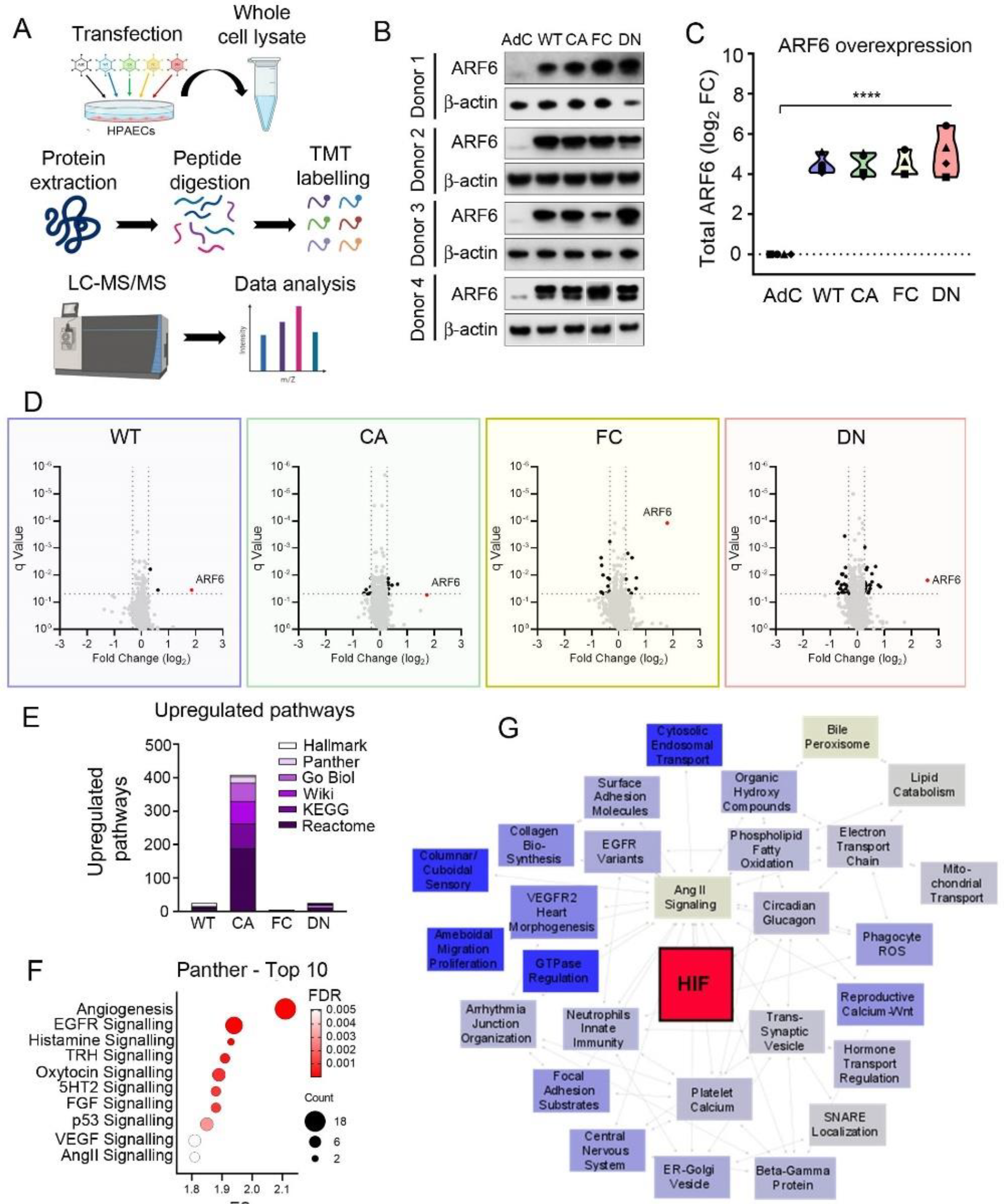
Proteomic analysis of HPAECs overexpressing ARF6 activity mutants. (**A**) Proteomics workflow for HPAEC whole cell lysates using tandem mass tag (TMT) labelling followed by liquid chromatography with tandem mass spectrometry. (**B**) Western blots and (**C**) quantification of total ARF6 overexpression in HPAECs from four different donors (n=4). (**D**) Volcano plots displaying the protein changes according to proteomic analysis for the four different ARF6 mutants; black symbols indicate differentially expressed proteins with ARF6 highlighted in red. (**E**) Stacked bar chart showing the number of upregulated pathways induced by each ARF6 mutant, according to each bioinformatics database. (**F**) Bubble plot of the 10 most significantly upregulated Panther pathways by overexpression of CA-ARF6. (**G**) Enrichment map summarising all upregulated pathway changes induced by CA-ARF6, with ‘HIF Signalling’ highlighted in red.

Enrichment analysis on ranked lists of all identified proteins against a range of curated databases (GO, Hallmark, KEGG, Panther, Reactome, WikiPathways) revealed that CA-ARF6 induced the highest number of pathway changes (559 upregulated, 282 downregulated) compared with WT-ARF6 (54 upregulated, 238 downregulated), FC-ARF6 (46 upregulated, 300 downregulated) or DN-ARF6 (117 upregulated, 292 downregulated) (**Fig. 2E)**. CA-ARF6 specifically enriched pathways associated with hypoxic signalling, including HIF, growth factors (EGF, FGF, VEGF), vasoconstrictors (histamine, serotonin, ET-1, Ang II), regulators of angiogenesis and glycolysis (**Fig. 2F**) and (**Table S8**). An enrichment map integrating data from all the bioinformatics databases further highlighted the regulatory role of ARF6 in HIF signalling, in addition to its well documented role in the vesicle transport (7) (**Fig. 2G**). Employing multiple databases was crucial given that different pathway databases generate disparate results in the statistical enrichment analyses (12).

### ARF6 induces normoxic activation of HIF

HIF transcription factors are master-regulators of gene expression in PAH, with HIF-2 thought to play a causative role in the disease (13-15). HIF-2 activation by CA-ARF6 was confirmed by its nuclear translocation (log_2_ FC 2.19±1.10, comparison with controls), while HIF-1 remained relatively unaffected (**Fig. 3A-C**). HIF activation was also confirmed by increased expression of luciferase reporter (log_2_ FC 0.52±0.17 change, comparison with controls) (**Fig. 3D)** and increased expression of HIF-regulated genes, including *PGK1* (log_2_ FC 0.46±0.24), *NOX4* (log_2_ FC 0.88±0.36), *RAB33A* (log_2_ FC 1.01±0.51) and *SLC2A1* (log_2_ FC 0.50±0.30) (**Fig. 3E**).

**Figure 3.**
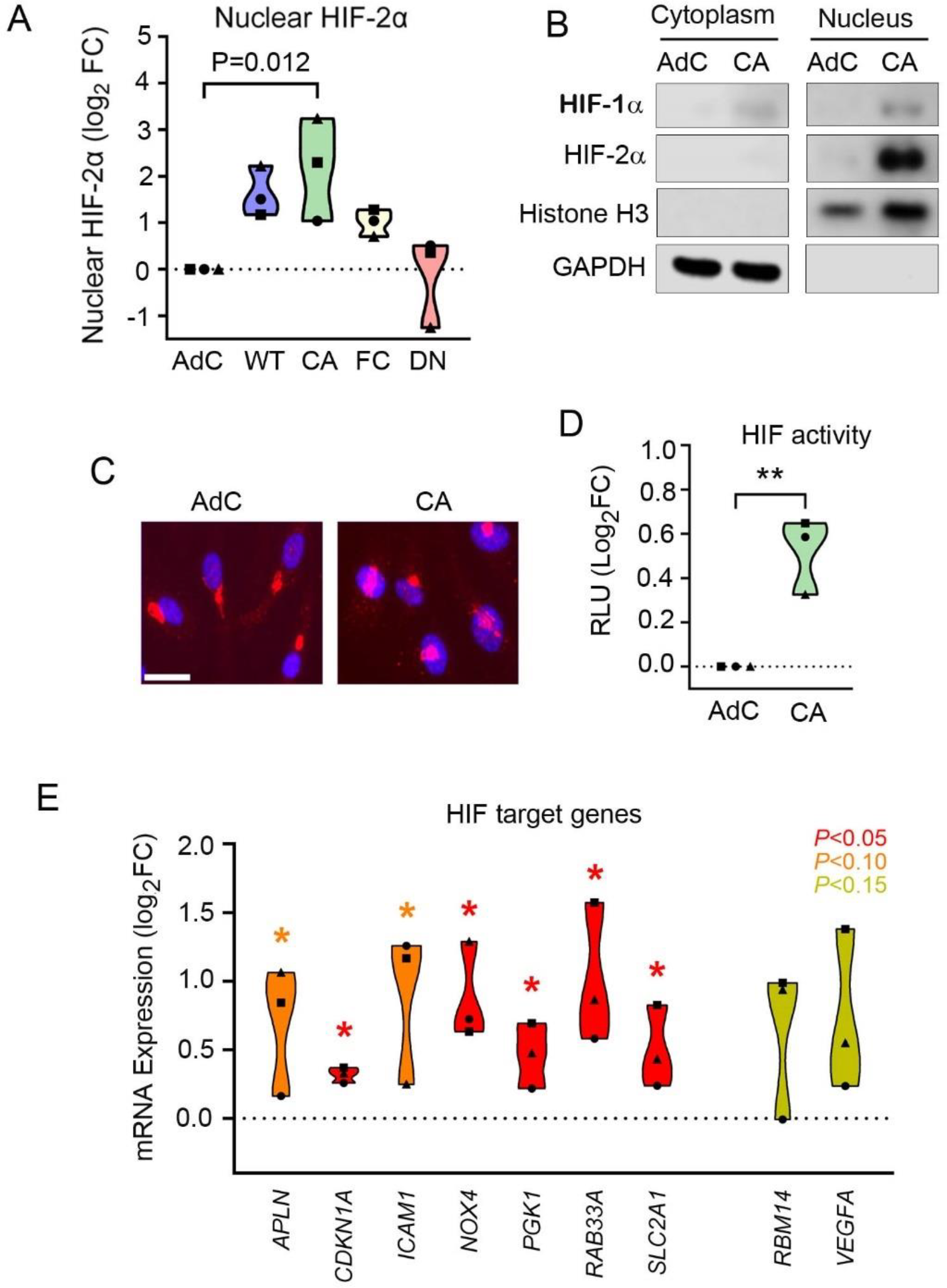
Active ARF6 induces HIF activity in HPAECs. (**A**) Quantification of HIF-2α in nuclear fractions of HPAECs overexpressing ARF6 mutants, as indicated; western blotting; n=3. (**B**) Representative western blots showing cytoplasmic and nuclear localization of HIF-1α and HIF-2α. (**C**) Cellular distribution of HIF-2α (red) merged with nuclear stain (DAPI, blue), immunofluorescence; Bar = 20μm. (**D**) HIF activity measured in luciferase reporter assay in U2OS-HRE-Luc cells (RLU = relative luminescence units; n=3 independent experiments). (E) Expression of selected HIF target genes by RT-qPCR (n=3).

### Effect of ARF6 activity on endothelial barrier function and proliferation *in vitro*

CA-ARF6 enhanced expression of important structural components of endothelial adherens and tight junctions, including CDH5 (VE-Cadherin), PECAM-1 (CD31) and JAM3 as well as adhesion receptors for leukocyte recruitment, ICAM-1 and ICAM2 (**Fig. 4A**). CA-ARF6 significantly upregulated junctional signalling pathways and altered endothelial barrier function (**Fig. 4B, C**), while other mutants had no effect. Increased expression of VE-cadherin was accompanied by its enhanced internalization (**Supplementary Figure S4**), consistent with the documented role of ARF6 in the regulation membrane trafficking of this protein (16).

**Figure 4.**
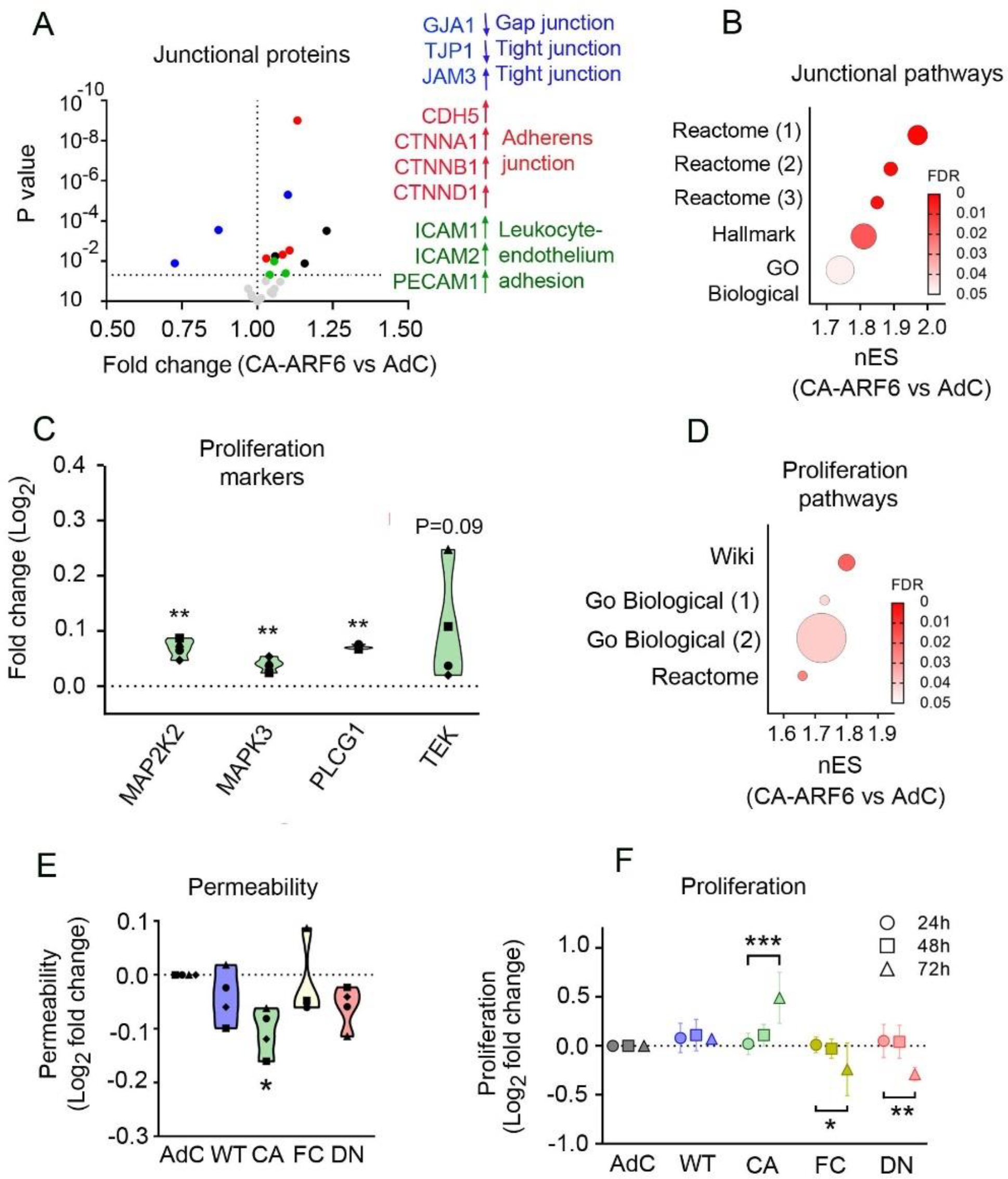
Active ARF6 alters endothelial junctions and increases endothelial proliferation. (**A**) Volcano plot of selected junctional proteins from proteomic analysis of CA-ARF6. (**B**) Bubble plot showing enrichment of pathways referencing ‘Junctions’ from different bioinformatics databases by CA-ARF6. (**C**) Bar chart of selected proliferation marker proteins from proteomic analysis of CA-ARF6. (**D**) Bubble plot showing enrichment of pathways referencing ‘Proliferation’ from different bioinformatics databases by CA-ARF6. (E) FITC-dextran permeability assay (n=4). (F) CyQUANT cell proliferation assay (n=3-4). In (**E**) *P<0.05, comparison with AdC; In (**F**) *P<0.05, **P<0.01, ***P<0.001, comparisons, as indicated; one-way ANOVA with Tukey post-test.

CA-ARF6 increased expression of numerous cell proliferation markers, including *MAP2K2* (MEK2), *MAPK3* (ERK1), *PLCG1* (PLCγ1) and *TEK* (TIE2) (**Fig. 4D**), which resulted in enrichment of proliferation-related pathways (**Fig. 4E**) and a time-dependent enhancement of HPAEC proliferation, particularly evident after a prolonged time of overexpression (72h) (**Fig. 4F**). Cell apoptosis was not affected by ARF6 mutants (**Supplementary Figure S5**). CA-ARF6 overexpression did not significantly alter expression or activity of the closely related ARF1 or other ARF family proteins (**Supplementary Figure S6**).

In summary, we show that sustained activation of ARF6 triggers normoxic activation of HIF-2α and, to a lesser extent, HIF-1 and that this effect is accompanied by endothelial junctional remodelling and increased endothelial cell proliferation.

### ARF6 regulates hypoxia-induced HIF activation in HPAECs

Hypoxia is a key contributor to vascular remodelling in PAH (17). HPAECs exposed to hypoxia (1% O_2_) showed a time-dependent increase in ARF6 activity, which peaked at 16 hours and remained elevated between 16-24 hours of hypoxic exposure (**Fig. 5A-B)**, while ARF6 protein expression remained relatively unaffected (**Supplementary Figure S7**).

**Figure 5.**
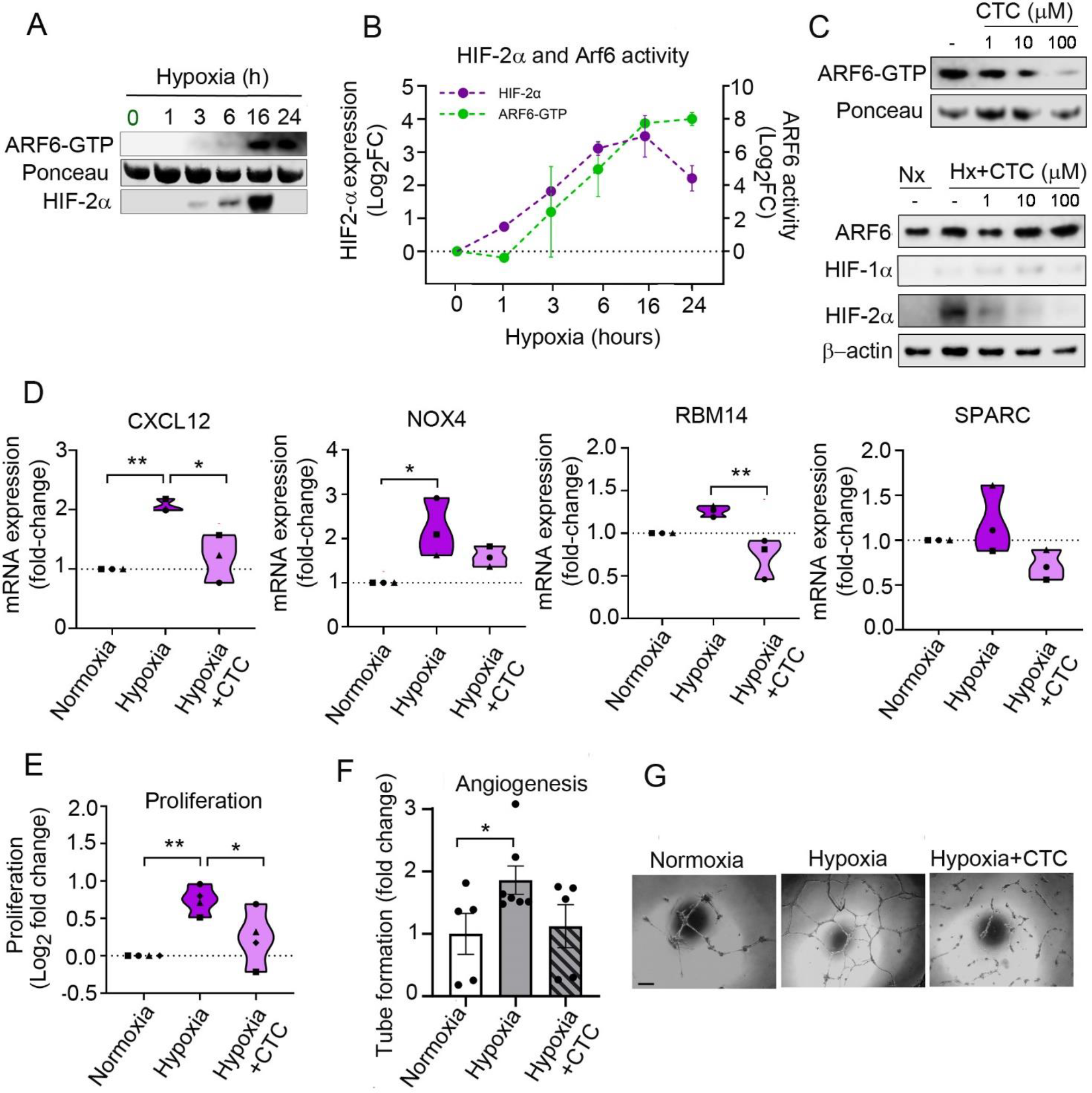
ARF6 Inhibitor Chlortetracycline (CTC) Modulates HIF Activation in Hypoxia. (**A**) Representative Western blots and (**B**) quantification of active ARF6-GTP and HIF-2α during hypoxia (1% O_2_) time course (n=4). (**C**) Representative western blots showing ARF6-GTP, total ARF6, HIF-1α and HIF-2α following treatment with increasing concentrations of chlortetracycline (CTC) in hypoxia (Hx, 24h), compared with normoxic control (Nx). (**D**) Expression of selected HIF target genes; qPCR, fold change of control; n=3-4. (**E**) CyQUANT cell proliferation assay; n=4; log2 fold change of control. (**F, G**) graph and corresponding representative images of endothelial tube formation in vitro. HPAECs were cultured in serum- and growth factors-reduced medium (0.5% FBS) and were treated, as indicated. n=5-6. *P<0.05; **P<0.01, comparisons, as indicated; one-way ANOVA with Tukey’s multiple comparisons test. In (G) Bar=100μm

The temporal activation profile of ARF6 peaked after 16 hours of hypoxia, which mirrored the time course of HIF-2α accumulation in HPAECs **(Fig. 5A, B**). HIF1-α activation in HPAECs, reported to reach maximum within 2-6 hours of the onset of hypoxia (18), did not correspond with changes in ARF6 activity.

The mechanism of ARF6-induced HIF-2α activation is likely to be complex, given the high number of upstream regulators and downstream effectors of both proteins (7, 17). Our proteomics data links ARF6 activation with epidermal growth factor receptor (EGFR) activity as well as other pathways such as PI3K-Akt-CK1δ and Ras-MAPK/ERK (**Supplementary Figure S8 a-e**), implicated in the regulation of HIF (19-22). In support of potential role of Akt/PI3K, we observed a significant CA-ARF6-induced activation of Akt (**Supplementary Figure S8 f-h**). In addition, the Akt inhibitor, MK-2206 (1μM)(23) and the PI3K inhibitor, LY294002 (50μM) (24) reduced ARF6-induced HIF-2 expression (**Supplementary Figure S8 h**).

To further verify the role of ARF6 in the regulation of HIF activity, we tested the effects of the newly identified, ARF6-specific inhibitor chlortetracycline (CTC) (25). CTC reduced protein expression of HIF-1α and HIF-2α in a concentration-dependent manner, reduced expression of HIF-1α and HIF-2α target genes in hypoxic HPAECs (**Fig. 5C-D**) and inhibited hypoxia-induced endothelial cell proliferation and angiogenesis **(Fig. 5E, F, G**). These findings demonstrate for the first time that ARF6 acts as a regulator of HIF-dependent endothelial responses to hypoxia *in vitro*.

### ARF6 regulates HIF signalling at the onset of hypoxia *in vivo*

To further verify our results and to test ARF6 role in the regulation of hypoxia-induced HIF signalling *in vivo*, mice were exposed to acute hypoxia (10% O_2;_ 24 hours), with one group receiving CTC (ip, 77.3mg/kg), prior to hypoxic exposure (**Fig. 6A**).

**Figure 6.**
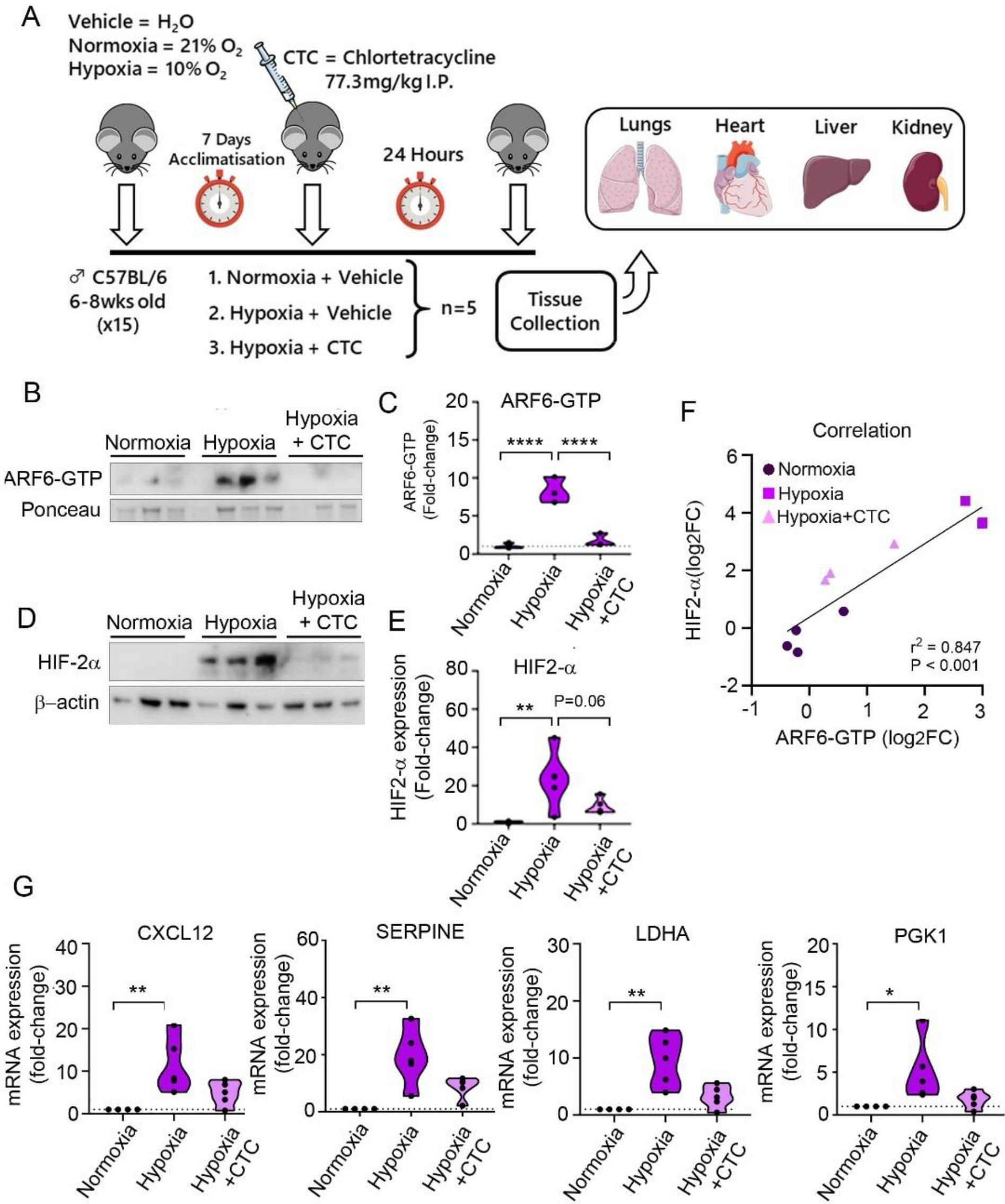
CTC Inhibits ARF6-Mediated HIF-2α activation in vivo. (**A**) Experimental outline: CTC (ip, 77.3mg/kg) or vehicle (ip, H_2_O) were administered to control mice or mice kept in a normobaric hypoxic chamber (10% O_2_) for 24 hours, as indicated. Lung, heart, kidney and liver tissues were collected from all animals. (**B, C**) Representative western blots and quantification of active ARF6 and (**D, E**) western blots and quantification of HIF-2α in mouse lung tissue lysates (n=3-4). (**F**) Pearson correlation between fold-changes in ARF6-GTP and HIF-2α based on western blotting analysis. (**G**) Expression of selected HIF target genes by RT-qPCR (n=4-5). *P<0.05; **P<0.01, comparisons, as indicated. one-way ANOVA with Tukey post-test.

Hypoxia induced a significant increase in ARF6 activity in mouse lung tissues, which was reduced to normoxic levels by CTC (**Fig. 6B-C**). Accumulation of HIF-2α in hypoxia was reduced by CTC treatment (**Fig. 6B-C**) and changes in HIF-2α activity positively correlated with changes in ARF6 activity (**Fig. 6D**). CTC markedly attenuated expression of HIF2 in hypoxic lung and heart but not in kidney (**Fig. 6E; Supplementary Figure S9**), suggesting that its actions are tissue-specific. Lung expression of HIF-1 was practically undetectable, but heart and kidney showed high expression of HIF-1, which was reduced by CTC (**Supplementary Figure S9**). These results reaffirm the role of ARF6 as an upstream regulator of HIF-2 and show that CTC, a novel and clinically available ARF6 inhibitor, reduces hypoxia-induced HIF signalling *in vivo*.

### ARF6 inhibition attenuates development of chronic hypoxia PH in mice

HIF signalling plays a key role in the initiation and development of chronic hypoxia-induced PH and here we studied the effects of CTC, administered in drinking water (40mg/kg/day) to mice exposed to 14 days of hypoxia (10% O_2_) (**Fig. 7A**). CTC significantly reduced right ventricular hypertrophy (**Fig. 7B**) right ventricular systolic pressure (**Fig. 7C**) and significantly reduced muscularization of small intrapulmonary arteries (**Fig. 7D, E**).

**Figure 7.**
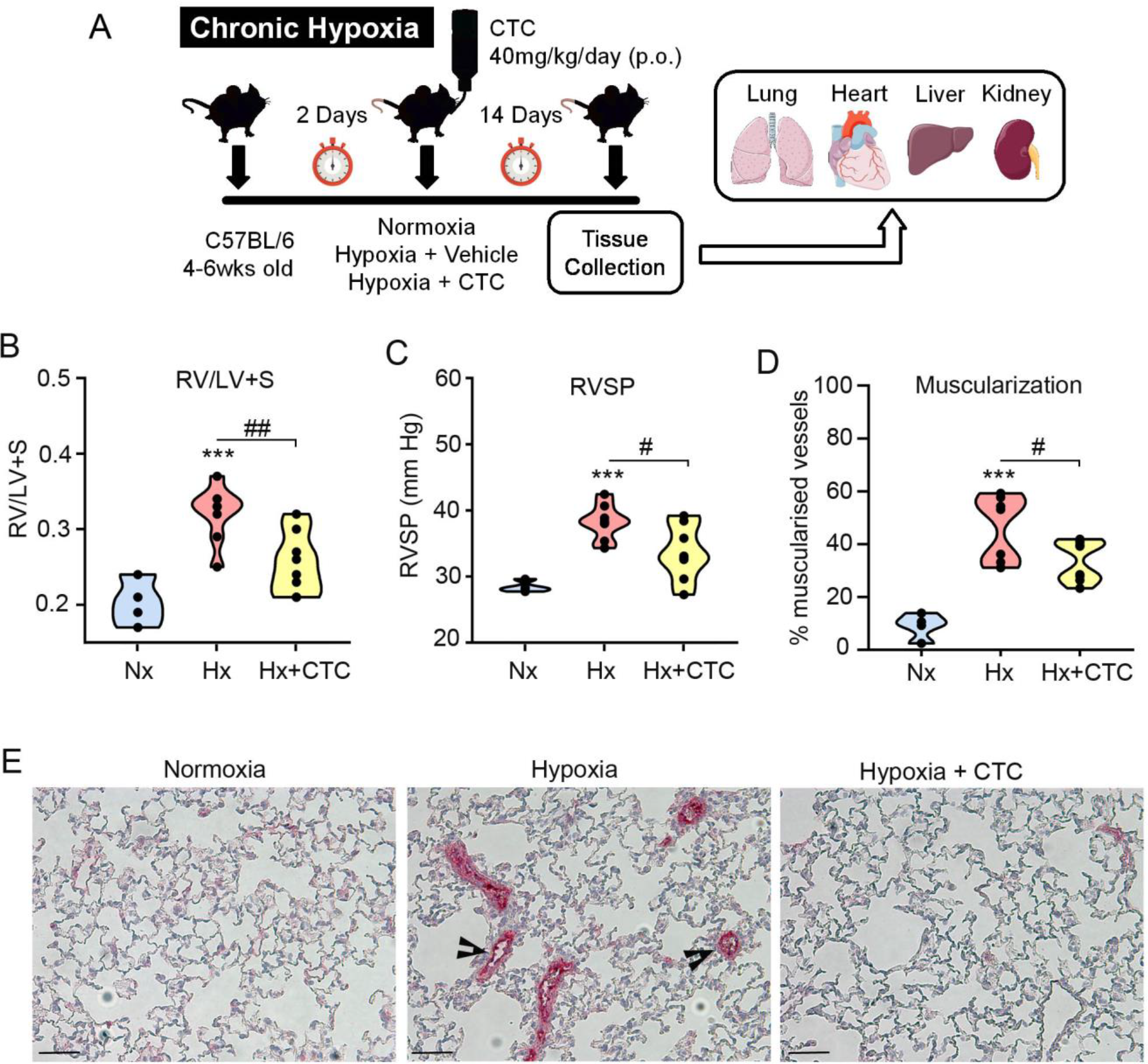
CTC prevents chronic hypoxia-induced PH in mice. (**A**) Experimental layout: C57/BL mice were left in normoxia (Nx) or were placed under hypoxia (Hx; 10% O_2_) for 2 weeks. CTC were administered to mice at the onset of hypoxia in drinking water (40mg/kg/day) (Hx+ CTC); (**B**) RV/LV+S; (**C**) RVSP; (**D**) Muscularization expressed as % of α-SMA-positive vessels compared with the total number of vessels with diameter <25μm. (**E**) Representative images of lung sections from different study groups, as indicated. Alkaline phosphatase immunostaining, α-SMA shown in pink. Arrowheads point to remodelled vessels. Bar = 40μm. In (**B-D**) ***P<0.001, comparison with normoxic controls; #P<0.05, comparison between Hx and Hx+CTC, n=4 in Nx and n=8 in Hx and Hx+CTC

### ARF6 signalling in PAH

To gain a further insight into the role in PAH, we identified ARF6 targets in publicly available PAH proteomic and transcriptomic PAH databases from PAH lung (26), blood-derived endothelial cells (27), HPAECs (28) and plasma (29) (**Fig. 8A**).

**Figure 8.**
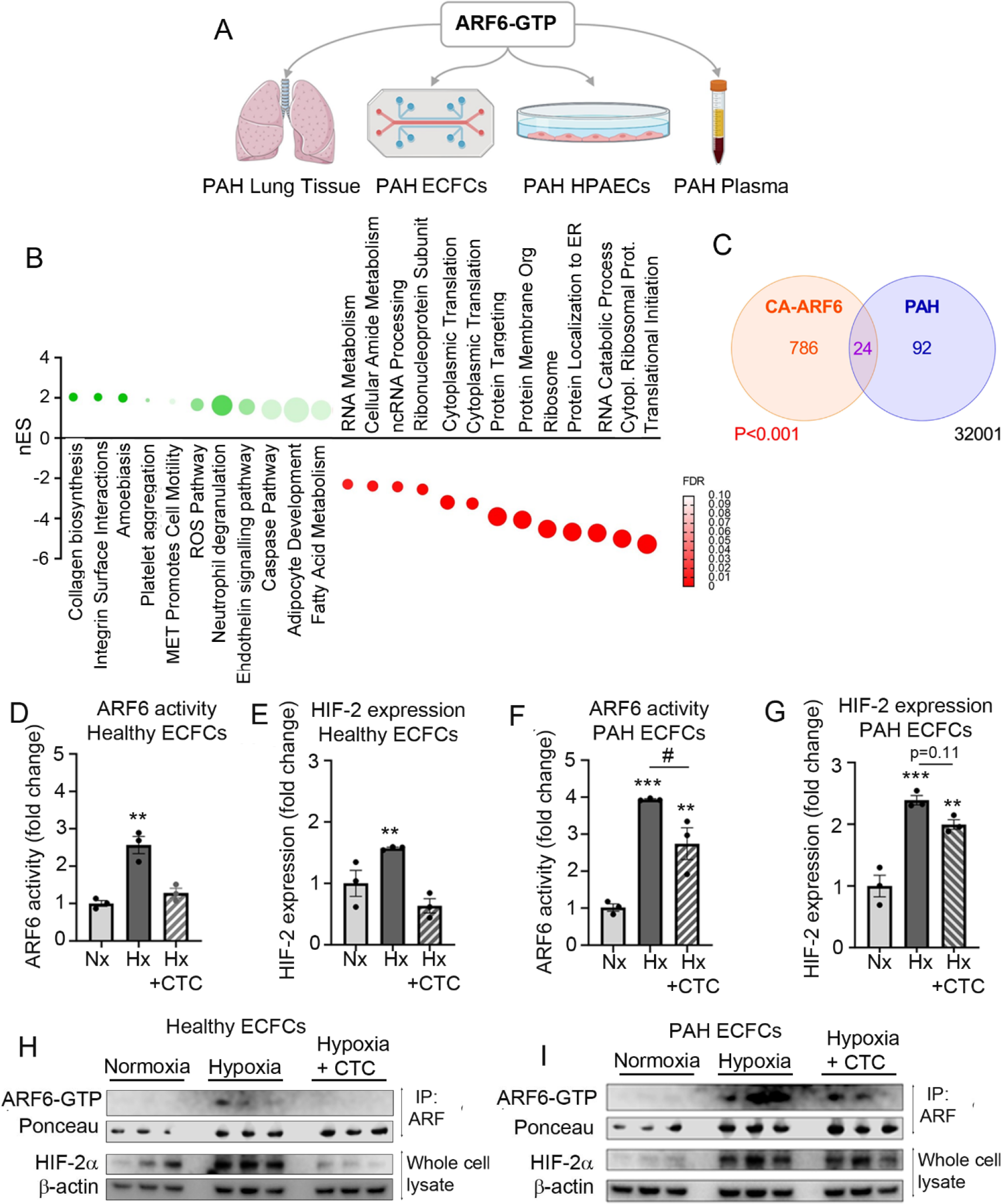
CTC restores healthy phenotype in PAH endothelial cells. (**A**) Schematic outline of comparative analysis between CA-ARF6 and other PAH databases. (**B**) Bubble plot of all shared upregulated and downregulated pathways in CA-ARF6 and PAH datasets. (**C**) Venn diagram showing number of pathways affected by CA-ARF6 and PAH. Statistical differences in proportions were analysed by Fisher’s Exact Test. (**D, E**) Graphs showing quantification of ARF6-GTP and HIF-2α in healthy and **(F, G**) PAH endothelial colony forming cells (ECFCs).(**H**) Representative western blots corresponding to (D, E) and (I) western blots corresponding to (F, G). In (D-G) **P<0.01, **P<0.001, one-way ANOVA with Tukey post-test, n=3.

ARF6 and PAH shared 24 pathways, including ‘response to hypoxia’, ‘endothelin signalling’, ‘fatty acid metabolism’, ‘reactive oxygen species production’, ‘collagen biosynthesis’ and ‘RNA processing’ (**Fig. 8B**) and this association was significant (P<0.01) (**Fig. 8C**).

Endothelial colony-forming cells (ECFCs) are commonly used as accessible surrogates for studying endothelial function in PAH (30). Healthy ECFCs responded to hypoxia in a similar manner to HPAECs, showing a CTC-responsive activation of ARF6 and HIF-2 (**Fig. 8D, E**). No changes to ARF1 activity or HIF-1 expression were observed (**Supplementary Figure S10**).

PAH ECFCs displayed a markedly enhanced response to hypoxia, manifested by a 4-fold increase in ARF6 activity and a 2.5-fold increase in HIF-2 levels, compared with a 2-fold and 1.5-fold change in respective healthy controls (**Fig. 8F, G**). Treatment of PAH cells with CTC reduced ARF6 activity and HIF-2α expression (**Fig. 8F, G**) and its selected target genes implicated in endothelial dysfunction in PAH, including NOX4 (NADPH oxidase 4), RBM14 (RNA binding motif protein 14) and SPARC (osteonectin) (**Supplementary Figure S10**).

## DISCUSSION

This study is first to describe ARF6-induced changes in pulmonary endothelial proteome and reveal a novel link between ARF6 with HIF-2 signalling, important in the pathogenesis of PAH.

ARF6 regulates numerous processes of critical importance in PAH, such as endocytotic trafficking, actin polymerization, cell migration, proliferation, angiogenesis and energy metabolism (5, 8). ARF6 has been implicated in the regulation of membrane trafficking of bone morphogenetic protein receptor 2 (BMPRII), the protein critically involved in vascular remodelling in PAH (4). The contribution of Arf6 is likely to extend beyond the regulation of BMPRII but mechanisms are not well characterised, as ARF6 proteome and interactome are largely unknown and current pharmacological activators and inhibitors are indirect and therefore non-specific.

To better understand the mechanism of ARF6 actions, we overexpressed four different ARF6 activity mutants in HPAECs, where ARF6 either remained unaltered (WT), was permanently activated (CA), inactivated (DN) or showed a spontaneously higher rate of GTP exchange and therefore a higher activity than the WT protein (FC). The inclusion of FC was important, as CA and DN, which block the normal cycle of GTP binding, hydrolysis and GDP release, may exert similar effects (6, 10). The analysis of endothelial proteome in cells overexpressing ARF mutants revealed a novel link between ARF6 and HIF signalling, and confirmed its associations with endosomal trafficking, angiogenesis and proliferation pathways. Experimental validation showed that active ARF6 increased expression and activity of HIF-2 in endothelial cells under normoxic and hypoxic conditions, *in vitro* and *in vivo*.

Given the complexity of ARF6 interactions, the mechanisms involved in HIF activation may not be easy to delineate. The most likely scenarios, based on the results of proteomic analysis and summarised in Figure S8a, implicate EGFR, PI3K-Akt and Ras-MAPK/ERK as upstream regulators of HIF, consistent with previous reports (19-22). Activation of Akt and inhibition of HIF-2 by Akt/PI3K inhibitors in CA-ARF6 overexpressing HPAECs argue for the involvement of Akt/PI3K pathway, previously implicated in HIFα transcription, stabilization, translation, and coactivator recruitment (31). Activation of Akt by ARF6 may be mediated by lipid-modifying enzymes. ARF6 activates phosphatidylinositol 4-phosphate 5-kinase (PtdIns4P5K) which synthesises phosphatidylinositol 4,5-bisphosphate (PtdIns(4,5)P(2)), required for the activation of Akt/PI3K (32), and phospholipase D1 which produces phosphatidic acid, known to activate HIF (5, 33). Whilst our data implicate ARF6 in the regulation of HIF-2 in HPAECs and ECFCs, we can not rule out its role in the regulation of HIF-1 in other types of cells or tissues. The precise mechanism by which ARF6 activates HIF, will require further investigation.

ARF6 activation has been linked with increased cell proliferation, migration, angiogenesis and inhibition of mitochondrial respiration (Warburg effect) (5, 34), processes which are also regulated by HIF-2 (35) and play a key role in PAH pathogenesis (13-15). We focused our investigation on the pulmonary endothelium, because of its key role in the initiation and progression of PAH. As a proof of concept and to verify the link between ARF6 and HIF-2 *in vivo*, the effects of ARF6 inhibition were studied in a pre-clinical model of chronic hypoxia-induced PH. Therapeutic effects of CTC, manifested by a significant reduction in RVSP, RVH and vascular remodelling, were accompanied by reduction in HIF-2 expression in lung and heart but not in systemic tissues, which suggests that CTC preferably affects organs most affected by the disease. Considering that ARF6 is expressed, to a varying degree, in all cells and tissues and has a broad spectrum of effectors, future studies should address the role of ARF6 signalling in other cell types. Regulation of inflammatory cell responses, critical in PAH, may be of particular interest (8, 36, 37).

CTC (also known as ‘aureomycin’) is a cell permeant first-generation antibiotic of the tetracyclin family, clinically approved for treatment of topical infections and widely used in veterinary medicine. CTC inhibits ARF6 GTP/GDP exchange by interacting with the Mg^2+^ present in the nucleotide binding site of ARF6 and, to a lesser extent, ARF1 (25). As ARF6 activity changes did not affect ARF1 expression or activity in our study, it is unlikely that ARF1 mediated any of the observed effects in HPAECs or ECFCs. However, we can not exclude a possibility that CTC may affect ARF1 in other cell types.

Unlike other ARF inhibitors, CTC is ARF6-specific, can be administered orally, is quickly absorbed from the gastrointestinal tract and undergoes minimal metabolic modifications. However, its potential clinical applications may be limited by gastrointestinal adverse effects and therefore more extensive studies will be required to evaluate its suitability as a drug candidate in PAH.

In conclusion, we show that ARF6 is a key mediator in HIF-2 signalling in pulmonary vascular endothelium. Protective effects of ARF6 inhibition in patient-derived cells and pre-clinical model of PAH re-enforce the view of ARF6 as a potential therapeutic target in PAH.

## METHODS

Detailed information can be found in Supplemental Materials and Methods.

### Cell Culture

Human pulmonary artery endothelial cells (HPAECs) from 6 different donors were purchased from Promocell (C-12241) and Lonza (CC-2530) and were cultured as previously described(38). HPAECs were used at passage 3-7. Donor information is provided in **Supplementary Table S1**.

### Cell treatments

Confluent HPAECs were treated with CTC (Cambridge Bioscience, C-2951) at a concentration of 100μM (unless otherwise stated), with H_2_O as a vehicle control. MK-2206 (Apexbio A3010-APE) and LY-294,002 (Sigma Aldrich, L9908) were used at specified concentrations with DMSO as a vehicle control. In some experiments HPAECs were exposed to hypoxia (1% O_2_) for 24-72h.

### Adenoviral gene transfer

HPAECs were infected with recombinant adenoviruses to overexpress haemagglutinin (HA-) tagged ARF6 mutants: wild-type (WT-ARF6), constitutively active ARF6Q67L(CA-ARF6), fast cycling ARF6T157A (FC-ARF6) and dominant negative ARF6T27N (DN-ARF6). Empty adenoviruses (Vector Biolabs; 1.0 x 10^12^pfu/mL) were used as adenoviral controls.

The ARF plasmids from Adgene (pcDNA3 HA-Arf6 #10834; pcDNA3 HA-Arf6 ActQ67L #10835 and pcDNA3 HA-Arf6 dnT27N #10831) were used to produce recombinant adenoviruses (Welgen, Inc, MA, USA).

Adenoviral infection was carried our as previously described (4). Experiments were performed 24 hours post-infection. ARF6 overexpression was confirmed by Western blotting, qPCR and immunocytochemistry.

### Proteomics – Sample Preparation

HPAECs overexpressing AdC, WT, CA, FC and DN ARF6 for 24 hours were lysed in RIPA buffer (EMD Millipore, 20-188) containing protease inhibitors. Protein quantification was determined using a bicinchoninic acid (BCA) assay kit (Thermo Fisher, 23227) according to manufacturer’s instructions. Following validation of robust and equal overexpression of mutant ARF6 protein by Western blotting, 10μg sample aliquots plus two additional aliquots containing 10μg of sample pool were used for proteomics analysis. Sample preparation procedure is described in Supplemental Materials and Methods.

### Proteomics – Tandem-Mass Tag Peptide Labelling

Peptide samples were TMT-11plex labelled according to manufacturer’s instructions (Thermo Fisher). Samples and pool were then grouped in equal amounts, ensuring randomisation, and dried using a speed vac (Thermo Fisher Scientific, Savant SPD131DDA). Grouped TMT samples were then reconstituted with 0.1% TFA in H_2_O, for peptide fractionation using high pH reversed-phase spin columns, following manufacturer’s instructions (Thermo Fisher).

### Proteomics – LC-MS/MS Analysis

The dried peptide samples for label-free and TMT analyses were reconstituted with 0.05% TFA in 2% ACN and separated by a nanoflow LC system (Dionex UltiMate 3000 RSLC nano). Samples were injected onto a nano-trap column (Acclaim® PepMap100 C18 Trap, 5mm x 300μm, 5μm, 100Å), at a flow rate of 25μL/min for 3 minutes, using 0.1% FA in H_2_O. The following nano-LC gradient was then run at 0.25μL/min to separate the peptides: 0–10min, 4–10% B; 10–75min, 10– 30% B; 75−80min, 30−40% B; 80–85min, 40–99% B; 85–89.8min, 99% B; 89.8– 90min, 99–4% B;90–120min, 4% B; where A = 0.1% FA in H2O, and B = 80% ACN, 0.1% FA in H_2_O. The nano column (EASY-Spray PepMap® RSLC C18, 2μm 100Å, 75μm x 50cm), set at 40°C was connected to an EASY-Spray ion source (Thermo Scientific). Spectra were collected from an Orbitrap mass analyser (Orbitrap Fusion™ Lumos Tribrid, Thermo Scientific) using full MS mode (resolution of 120,000 at 400m/z) over the mass-to-charge (m/z) range 375–1500. Data-dependent MS2 scan was performed using Quadrupole isolation in Top Speed mode using CID activation and ion trap detection in each full MS scan with dynamic exclusion enabled. TMT reporter ion quantification was conducted using the Multi-Notch MS3 method(39); the top 5 most abundant MS2 fragment ions with an MS isolation window (m/z) of 0.7 were isolated, following HCD activation and detection using the Orbitrap at a resolution of 60,000, generating an MS3 spectrum.

### Proteomics – MS Database Search and Analysis

Thermo Scientific Proteome Discoverer software (version 2.2.0.388) was used to search non-crosslinked raw data files against the human database (UniProtKB/Swiss-Prot version 2018_02, 20,400 protein entries) using Mascot (version 2.6.0, Matrix Science). The mass tolerance was set at 10ppm for precursor ions and 0.8Da for fragment ions. Trypsin was used as the digestion enzyme with up to two missed cleavages being allowed. Carbamidomethylation of 12 cysteines and oxidation of methionine residues were chosen as fixed and variable modifications, respectively. The in-built TMT 11-plex quantification method was assigned for detection of TMT labels. MS/MS-based peptide and protein identifications were validated with the following filters, a peptide probability of greater than 95.0% (as specified by the peptide prophet algorithm), a protein probability of greater than 99.0%, and at least two unique peptides per protein. TMT-acquired data was again normalised using the total peptide amount. Data was then further scaled using the control pooled sample abundance, correcting for technical variation between injections and TMT groups(40).

### Proteomics – Statistical Analysis

Proteomic data were filtered to keep only molecules with less than 70% missing values. The remaining missing values were imputed using k-nearest neighbour-imputation method with k equal to 10. The relative quantities of the molecules were normalised to the control (empty adenovirus; AdC) of the corresponding biological replicate to calculate the fold-change then scaled using log_2_ transformation to perform statistical testing. To generate volcano plots, log_2_ fold-changes for each protein were analysed by an unpaired Student’s t-Test then adjusted for multiple comparisons using the original FDR method of Benjamini and Hochberg with a q value of 5%. These q values were plotted against log_2_ fold-change, with thresholds of Q<0.05 and FC±0.2 indicated by dotted lines. Proteins which met both of these thresholds were deemed differentially expressed.

### Bioinformatics

Bioinformatics was performed using the functional enrichment analysis web tool WebGestalt (www.webgestalt.org). The basic parameters were as follows: Organism of Interest: *Homo Sapiens*; Method of Interest: GSEA (Gene Set Enrichment Analysis); Functional Database: GeneOntology (GO, Biological Process), Pathway (KEGG, Panther, Reactome, WikiPathway), Network (Kinase Site), Disease (DisGenet, GLAD4U, OMIM), Drug (DrugBank, GLAD4U), Phenotype (Human Phenotype Ontology), Community-Contributed (Hallmark50). The results obtained from DisGenet, DrugBank, GLAD4U, Human Phenotype Ontology and OMIM did not yield data relevant to PAH. Proteins in each treatment group (WT-ARF6, CA-ARF6, FC-ARF6, DN-ARF6) were assigned a ranking score calculated by fold-change multiplied by the log_10_-transformed non-adjusted *P* value. These scores were then inputted with the following advanced parameters: Minimum Number of Genes for a Category: 5; Maximum Number of Genes for a Category: 2000; Significance Level: FDR≤0.1; Number of Permutations: 1000; Enrichment Statistic: 1; Collapse Method: Mean. Such thresholds were used, along with no prior filtering, due to the low number of biological replicates (i.e., relatively low statistical power), in order to maximise the number of significant pathways identified, with the stipulation that pathways of interest would be validated extensively. Results are presented in bubble plots as normalised enrichment score (nES), FDR (q value) and count (number of proteins).

All upregulated pathways identified by six bioinformatics databases (GO Biological Process, Hallmark, KEGG, Panther, Reactome, WikiPathway) induced by CA-ARF6 were integrated into an Enrichment Map using Cytoscape software (41, 42). The Enrichment Map and AutoAnnotate applications were used to create a summary network (excluding unclustered nodes) colour-coded by heat map according to the neighborhood connectivity of the pathway clusters.

### Western Blotting

Western blotting procedures with lists of primary and secondary antibodies are provided in Supplemental materials and Methods.

### ARF6 Activity Pull-Down Assay

Active, GTP-bound ARF6 was detected in cell lysates using the ARF6 Activation Assay Biochem Kit™ (Cytoskeleton, BK-033) according to manufacturer’s instructions.

### RT-qPCR

Protocols for RT-qPCR, with lists of human and mouse primers are provided in Supplemental Materials and Methods.

### Immuno- and Affinity staining

To stain ARF6 and F-actin in cells, HPAECs were fixed in 4% paraformaldehyde in PBS, permeabilised in 0.1% Triton X-100 in PBS and blocked in 5% bovine serum albumin (Sigma) in PBS. Fixed and permeabilized HPAECs were then incubated with mouse monoclonal anti-human ARF6 antibody (Santa Cruz; # sc7971, 1:100) or VE-Cadherin-AlexaFluor 488 (eBioscience; 16B1, # 53-1449-42; 1:100) and then FITC-Goat Anti-Mouse IgG (Jackson ImmunoResearch Inc., # 115-095-003; 1:200). To visualise F-actin, TRITC-phalloidin (Sigma, P1951) was added to the secondary antibody solution at the final concentration of 1mg/L. Cells were examined under the fluorescent confocal microscope (Leica, TCS SP5, Leica Biosystems).

### Luciferase Reporter Assay

NFκB activity was measured in luciferase reporter assay in the GlomaxTM luminometer(38).

### Tube formation Assay

HPAECs were seeded at the density of 10,000 cells/well in a pre-prepared 96-well plate containing 30μl of growth-factor reduced Matrigel (Merck, # CLS354230-1EA) per well (38). The cells were then cultured in growth factor-free, serum-reduced EGM2 medium (0.5% FBS) under normoxic or hypoxic conditions for 18 hours. Tube length was scored with Image J v. 2.3 (with Angiogenesis analysis plugin)

### Animal Studies

All studies were conducted in accordance with UK Home Office Animals (Scientific Procedures) Act 1986. All animals were randomly allocated to groups, and personnel involved in data collection and analysis (haemodynamics and histopathologic measurements) were blinded to the treatment status of each animal. Only weight-and age-matched males were included for experimentation as, in contrast to the human clinical studies, most animal studies have shown that female sex and estrogen supplementation have a protective effect against hypoxia-induced PH in rodents (43).

In the acute hypoxia study (24 hours), 10-12 weeks old C57/BL male mice (20 g; Charles River, UK) (n=15) were allowed to acclimatise in the facility for 7 days and then divided into three treatment groups (n=5/group): normoxia + vehicle (water), hypoxia + vehicle, hypoxia + CTC. The mice were left in normoxia or were placed in a normobaric hypoxic chamber (10% O_2_) for 24 hours. CTC was administered by intraperitoneal injection at 77.3mg/kg prior to hypoxic exposure.

In the chronic hypoxia-induced pulmonary hypertension study, 10-12 weeks old C57/BL male mice (20 g; Charles River, UK) (n=20) were allowed to acclimatise in the facility for 2 days and then were divided into three treatment groups: normoxia (n=4), hypoxia + vehicle (n=8), hypoxia + CTC (n=8). The mice were kept in normoxia or in a normobaric hypoxic chamber (10% O_2_) for 14 days. CTC was administered via drinking water at 40mg/kg/day, with water consumption and mice weight monitored every other day throughout the experiment. This is a dose recommended in veterinary practice (http://www.kepro.nl/wp-content/uploads/2014/06/Chlortetracycline-hydrochloride_0113_ENG-v3.pdf)

At 2 weeks mice were anaesthetised by intraperitoneal injection of Ketamine/Dormitor (75mg/kg + 1mg/kg). Development of PH was verified as described previously(38). Briefly, right ventricular systolic pressure (RVSP) was measured via direct cardiac puncture using a closed-chest technique in the spontaneously breathing, anesthetized animal. The animals were then euthanized, the hearts were removed, and the individual ventricular chambers were weighed, right ventricular hypertrophy was assessed as the right ventricle to left ventricle/septum ratio (RV/LV+S). The right lungs were snap-frozen in liquid nitrogen and stored at -80°C for biochemical measurements or placed in RNAlater® RNA Stabilization Solution for RNA isolation. The left lungs were fixed by inflation with 10% formalin, embedded in paraffin, and sectioned for histology. Transverse formalin-fixed lung sections were stained with hematoxylin and eosin (H&E) for routine examination or Verhoeff’s van Gieson stain (EVG) to visualise elastic lamina. Muscularised and partially muscularised vessels were visualised by immunostaining of sections with mouse monoclonal anti-α muscle actin (α-SMA) and VECTASTAIN® ABC-AP Kit, Alkaline Phosphatase (Standard) (Vector Laboratories #AK-5000).(44)

Muscularization of small intrapulmonary arteries was determined by counting all muscularised vessels (showing positive staining for αSMA) with a diameter smaller than 25μm in each section, and expressed as a % of all (muscularised + non-muscularised) vessels. All samples were scored simultaneously and blinded to the study condition.

### Late outgrowth Endothelial Colony Forming Cells (ECFCs)

Endothelial colony forming cells were derived from peripheral blood samples and characterised, as previously described (4, 27, 38). All ECFCs were used between passages 3-6. Demographic and clinical features of healthy subjects and PAH patients are provided in Supplemental Materials and Methods.

### Study approval

Venous blood samples were obtained with local ethics committee approval and informed written consent from healthy volunteers and idiopathic PAH patients. Participants were identified by number. Animal studies were conducted in accordance with UK Home Office Animals (Scientific Procedures) Act 1986.

### Apoptosis Assay

Apoptosis was measured using Cell MeterTM Caspase 3/7 Activity Apoptosis Assay Kit (AAT Bioquest, ABD-22796). Fluorescence intensity was analysed in GlomaxTM luminometer at Ex/Em= 490/525 nm.

### Permeability Assay

Changes in endothelial barrier function in HPAECs grown in Transwell-Clear filters were carried out as previously described(38).

### Proliferation Assay

HPAEC proliferation was studied in CyQUANT™ Cell Proliferation Assay (Thermofisher, # C7026), according to the manufacturer’s protocol.

### Receptor Internalisation Assay

Internalization of VE-cadhering in HPAECs overexpressing ARF6 mutants was carried out 24 hours post-infection. The method was adapted from (45).

### Statistical Analysis & Data Presentation

All graphs were analysed and plotted using Graphpad Prism software (Graphpad Software Inc, CA, USA). Statistical comparisons between three or more groups were performed with ordinary one-way or two-way ANOVA and Dunnett’s multiple comparisons test. Statistical comparisons between two groups were performed using unpaired Students t-test. Gaussian distributions and equal standard deviations between groups were confirmed prior to statistical tests. Violin plots were used to present data with appropriate P values or asterisks where *P<0.05, **P<0.005 and ***P<0.0005.

## ACKNOWLEDGEMENTS

This research was supported by the British Heart Foundation project grant PG/19/19/34286. BHF Imperial Centre of Research Excellence grant RE/18/4/34215. We thank the staff of the NIHR Imperial Clinical Research Facility, Hammersmith Hospital (London UK).

