## Supplement for "ARF6 as a novel activator of HIF-2α in pulmonary arterial hypertension"

**SUPPLEMENTAL MATERIALS AND METHODS    Fellows et al.**

**Cell Culture**

Human pulmonary artery endothelial cells (HPAECs) were purchased from Promocell (C-12241) and Lonza (CC-2530) and cultured in endothelial cell medium containing supplements (Promocell C-22111) and Pen/Strep (Gibco, 15140-122; 100U/ml) in a humidified incubator at 37°C containing 5% $CO_2$. Cells were split in an approximately 1:3 ratio using trypsin/EDTA (Gibco 25300-062). All flasks and plates were coated with 10μg/ml fibronectin (Sigma Aldrich, F1141) diluted in PBS (Sigma Aldrich D8537) prior to cell seeding. HPAECs were used at passage 3-7 and donor information is listed below.

| Assigned | Company | Catalogue # | Lot Number | Age | Sex |
|---|---|---|---|---|---|
| Donor 1 | Promocell | C-12241 | 431Z031 | 23 | Female |
| Donor 2* | Promocell | C-12241 | 433Z034 | 34 | Female |
| Donor 3# | Promocell | C-12241 | 458Z016.12 | 51 | Female |
| Donor 4 | Promocell | C-12241 | 451Z031.14 | 68 | Male |
| Donor 5* | Promocell | C-12241 | 433Z034.2 | 34 | Female |
| Donor 6# | Promocell | C-12241 | 458Z016.13 | 51 | Female |
| Donor 7 | Lonza | CC-2530 | 18TL155113 | 42 | Male |
| Donor 8 | ScienCell | 3100 | 9647 | - | - |

**Table S1.** HPAEC Donor Information (*, # Isolated from the same individual).

**Cell treatments.**

Confluent HPAECs were treated with CTC (Cambridge Bioscience, C-2951) at a concentration of 100μM (unless otherwise stated), with $H_2O$ as a vehicle control. MK-2206 (Apexbio A3010-APE) and LY-294,002 (Sigma Aldrich, L9908) were used at specified concentrations with DMSO as a vehicle control. In some experiments HPAECs were exposed to hypoxia (1% $O_2$) for 24-72h.

**Adenoviral gene transfer**

HPAECs (80-90% confluent) were infected with recombinant adenoviruses to overexpress haemagglutinin (HA-) tagged ARF6 mutants: wild-type (WT-ARF6), constitutively active ARF6Q67L(CA-ARF6), fast cycling ARF6T157A (FC-ARF6) and dominant negative ARF6T27N (DN-ARF6). Empty adenoviruses (Vector Biolabs; 1.0 x $10^{12}$pfu/mL) were used as adenoviral controls.

The ARF plasmids from Adgene (pcDNA3 HA-Arf6 #10834; pcDNA3 HA-Arf6 ActQ67L #10835 and pcDNA3 HA-Arf6 dnT27N #10831) were used to produce recombinant adenoviruses (Welgen, Inc, MA, USA). Briefly, the ARF6-HA, ARF6-ActQ67L-HA and ARF6-dnT27N-HA genes were subcloned into KpnI and XbaI restriction sites of pAd shuttle vector, pEntCMV. The purified pEntCMV-ARF6-HA, pEntCMV-ARF6-actQ67L-HA and pEntCMV-ARF6-dnT27N-HA ligated to a pAd-REP plasmid that contains the remaining adenovirus genome. The recombination pAd DNA were purified and digested with Pac1 and then transfected into 293 cells. The transfected 293 cells were grown at 37 °C with 5% CO2.

The low titer of virus was further amplified to $10^{12}$ vp/ml. The amplified adenoviruses were purified on 2 sequential caesium chloride gradients and then dialyzed with a buffer (PBS, 10% glycerol, pH 7.4) to reduce the salt concentration. Adenoviral infection was carried our as previously described (1). Experiments were performed 24 hours post-infection. ARF6 overexpression was confirmed by Western blotting, qPCR and immunocytochemistry.

**Proteomics – Sample Preparation**

HPAECs overexpressing AdC, WT, CA, FC and DN ARF6 for 24 hours were lysed in RIPA buffer (EMD Millipore, 20-188) containing protease inhibitor cocktail (Thermo Fisher, 78415) and phosphatase inhibitor cocktail (Sigma Aldrich, P0044). Information on donors used for proteomic analysis is provided in Supplementary Table S2.

| Donor | Age | Sex | Race | Passage |
|---|---|---|---|---|
| Donor 1 | 23 | Female | Caucasian | p4 |
| Donor 2 | 34 | Female | Caucasian | p6 |
| Donor 3 | 51 | Female | Caucasian | p7 |
| Donor 4 | 34 | Female | Caucasian | p5 |

**Table S2. Proteomics Study: HPAEC donor information.**

Data regarding age, sex and race of individuals from whom HPAECs were obtained (supplied by the manufacturer) and passage number at which whole cell lysates were generated for proteomics analysis.

Culture media were aspirated and cells were washed once with ice-cold PBS (Sigma Aldrich, D8537) before 100µl lysis buffer was added and then incubated on ice for 2-5 minutes. Lysates were collected by scraping (Sarstedt, 83.3951), triturated three times, transferred to 1.5ml protein Lo-bind eppendorfs (Sarstedt, 72.706.600) and centrifuged at 16,000g for 5 minutes to clear cellular debris. Lysates were stored at -80°C. Protein quantification was determined using a bicinchoninic acid (BCA) assay kit (Thermo Fisher, 23227) according to manufacturer's instructions. Following validation of robust and equal overexpression of mutant ARF6 protein by Western blotting, 10µg sample aliquots plus two additional aliquots containing 10µg of sample pool were used for proteomics analysis.

Protein samples were denatured by the addition of a final concentration of 6M urea and 2M thiourea and reduced by the addition of a final concentration of 10mM dithiothreitol (DTT) followed by incubation at 37°C for 1 hour at 240rpm. The samples were then cooled down to room temperature before being alkylated by the addition of a final concentration of 50mM iodoacetamide followed by incubation in the dark for 30 minutes. Pre-chilled (-20°C) acetone

(10x volume) was used to precipitate the samples overnight at -20°C. Samples were centrifuged at 14,000g for 40 minutes at 4°C and the supernatant subsequently discarded. Protein pellets were dried using a speed vac, resuspended in 0.1M TEAB buffer, pH 8.0, containing MS grade Trypsin/Lys-C (Promega Cooperation) (1:25 enzyme: protein) and digested overnight at 37°C, 240rpm. Peptides were then purified robotically, using C18 cartridges (Bravo AssayMAP, Agilent Technologies)(2).

**Western Blotting**

Cell lysates were prepared, quantified and stored as above or cell fractionation was performed with a kit (Cell Signaling Technology, 9038S) according to manufacturer's instructions. Lysates were thawed on ice and centrifuged at 16,000g for 5 minutes at 4°C then diluted with 4X reducing Laemmli buffer (Alfa Aesar, J60015). After, samples were gently vortexed, boiled at 98°C for 5 minutes in a heating block, cooled on ice for 2 minutes and centrifuged at 16,000g for 2 minutes. Samples (5-20µg) were loaded into 10-, 15- or 17-well NuPAGE 4-12% Bis-Tris polyacrylamide gels (Thermo Fisher, NP0335BOX, NP0336BOX, NP0329BOX) alongside a broad-range protein standard (Thermo Fisher, 26634) and separated by electrophoresis at 100-200V (Biorad, Powerpac 300) for 1-2 hours using SDS running buffer (Invitrogen, NP0002-02), then washed briefly in $H_2O$ and transferred to PVDF membranes (Biorad, 1704156) using the Transblot® Turbo™ transfer system (Biorad, 1704150). Next, membranes were washed in $H_2O$, blocked in milk (EMD Millipore, 1.15363.0500; 5% in TBST) for 1 hour and incubated at 4°C for a minimum of 16 hours in primary antibody, diluted in 5% BSA (Sigma Aldrich, A9418) in TBST (Thermo Fisher, J77500). After, membranes were washed three times in TBST for 15 minutes each then incubated with secondary antibody (diluted in 5% BSA/TBST) for 1 hour followed by three washes in TBST for 10 minutes each. Imaging was performed using Immobilon® Forte HRP substrate (EMD Millipore, WBLUF0100) and a ChemiDoc system (Biorad, 731BR01754). To reprobe, membranes were washed twice in PBS for 2 minutes each, incubated with stripping buffer (Thermo Fisher, 21059) for 5-15 minutes, washed again in PBS before the protocol was repeated as above from the blocking step. Correct bands were identified by their molecular weight and quantified using the gel imaging tool on ImageJ. All values were normalised to β-actin as a loading control.
Lists of primary and secondary antibodies used for western blotting are provided in **Table S2** and **Table S3** below.

**Table S3.** Primary antibodies (Western Blotting)

| Target | Host | Dilution | Company | Catalogue No. |
|---|---|---|---|---|
| ARF1 | Mouse | 1:50 | Santa Cruz Biotechnology | sc-53168 |
| ARF1-GTP | Mouse | 1:100 | Cytoskeleton | ARF01 |
| ARF6 | Mouse | 1:50 | Santa Cruz Biotechnology | sc-7971 |
| ARF6-GTP | Mouse | 1:50 | Cytoskeleton | ARF06 |
| Akt | Rabbit | 1:1000 | Cell Signaling Technology | 9272 |
| p-Akt (S473) | Rabbit | 1:500 | Cell Signaling Technology | 9271 |
| β-actin | Mouse | 1:100 or 1:500 | Santa Cruz Biotechnology | sc-47778 |
| GAPDH | Rabbit | 1:10,000 | Sigma Aldrich | G9545 |
| HIF-1α (Human) | Mouse | 1:250 | Enzo Life Sciences | OSA-602 |
| HIF-1α (Mouse) | Mouse | 1:500 | Thermo Fisher | MAI-16504 |

| | | | | |
|---|---|---|---|---|
| HIF-2α | Rabbit | 1:500 | Novus Bio | NB100-122 |
| Histone H3 | Rabbit | 1:1000 | Abcam | ab5103 |
| VE-Cadherin | Mouse | 1:100 | Santa Cruz Biotechnology | sc-9989 |

**Table S4.** Secondary antibodies (Western blotting)

| Target | Host | Conjugate | Dilution | Company | Catalogue No. |
|---|---|---|---|---|---|
| Rabbit IgG | Goat | HRP | 1:2000 | Sigma Aldrich | A6154-1ML |
| Mouse IgG | Sheep | HRP | 1:2000 | Sigma Aldrich | A6782-1ML |
| Mouse IgG | Sheep | HRP | 1:2000 | GE Healthcare | NA931 |
| Goat IgG | Rabbit | HRP | 1:2000 | Sigma Aldrich | A5420-1ML |

**ARF6 Activity Pull-Down Assay**

Active, GTP-bound ARF6 was detected in cell lysates using the ARF6 Activation Assay Biochem Kit$^{TM}$ (Cytoskeleton, BK-033) according to manufacturer's instructions. Briefly, lysates were obtained using the supplied lysis buffer and 30-60µg protein was diluted to 90µl with the same buffer. This was mixed with 10µl of GGA3-coated agarose beads for 1 hour at 4°C on a rotator. Beads were then centrifuged at 5,000g for 2 minutes, washed twice with wash buffer and resuspended in 1X sample buffer. GGA3-bound proteins were dissociated from the beads by boiling at 98°C for 2 minutes and then processed according to the same protocol as seen in 'Western Blotting' using supplied monoclonal antibodies for ARF1-GTP and ARF6-GTP. Recombinant ARF6 protein (Abcam, ab102025) was used as a positive control sample and membranes were stained with Ponceau S (Sigma Aldrich, P7170) immediately after transfer.

**RT-qPCR**

To obtain RNA, culture media was aspirated and 700µl TRIzol reagent (Life Technologies, 15596026) was added to cells, triturated gently three times, incubated at room temperature for 5 minutes, triturated three times again and transferred to RNase-free 1.5ml eppendorfs (Sarstedt, 72.706.600) and placed on ice. Samples were either processed immediately or stored at -80°C and later thawed on ice. Before going any further, all pipettes, surfaces and tip boxes (filter tips: Starlab, S1122-1830, S1120-8810, S1180-3810) were thoroughly sprayed with RNaseZAP (Invitrogen, AM9780). To isolate the RNA, samples were incubated at room temperature for 5 minutes, 140µl chloroform (Sigma Aldrich, C2432) was added and mixed well by inversion for 20 seconds, then samples were incubated for 3 minutes at room temperature before being replaced on ice. To precipitate the RNA, samples were centrifuged at 12,000g at 4°C for 15 minutes, 300µl of the aqueous phase was carefully collected into fresh eppendorfs, into which was added 4µl of glycogen (5mg/ml; Thermo Fisher AM9510) and 450µl of isopropanol (Thermo Fisher, 327272500) followed by gentle mixing by hand and 10 minutes incubation at 8°C. A pellet was formed following centrifugation at 12,000g at 4°C for 10 minutes which was subsequently washed in 1ml 75% EtOH (VWR, 437433T) in DEPC-treated $H_2O$ (Thermo Fisher, AM9960), gently vortexed and centrifuged again at 7,500g at 4°C for 5 minutes. Finally, the supernatant was aspirated and the pellet was allowed to dry for approximately 10 minutes on ice before being resuspended in 20-30µl RNase-free $H_2O$ (Invitrogen, 10977-035). RNA concentration and purity was determined with a NanoDrop 2000 spectrophotometer (RNA was deemed acceptable with an 260/280 ratio ≥1.80) then converted to cDNA by dilution of 0.5-1.0µg RNA to 16µl with $H_2O$ into which was added 4µl Luna RT Supermix (New England Biolabs, E3010). RT-PCR was performed using a SimpliAmp™ Thermocycler (Life Technologies) with the following protocol: 25°C for 2 minutes, 55°C 10 minutes, 95°C 1 minute, hold at 4°C. For qPCR, the concentration of cDNA was diluted to 5ng/µl with RNase-free $H_2O$, then 5ng cDNA was combined with a mixture of $H_2O$ (1.75µl), forward and reverse primers (0.35µl) and Luna Universal qPCR Mastermix (New England Biolabs, M3003E) (3.5µl) in 384-well plates (Starlab, E1042-3840). Real-time, quantitative PCR was performed using a QuantStudio 6 system (Thermo Fisher) with the following steps: Hold: 95°C – 1 min, PCR: 95°C – 15s, 62°C – 30s (40 cycles), Melt: 60°C – 1 min, then up to 95°C (0.05°C/s). Two housekeeping genes (Mouse: *B2m*, *Ywhaz*; Human: *TBP*, *YWHAZ*) were run in parallel and the most stable expression was used to calculate fold-changes for genes of interest according to the $2^{-\Delta\Delta Ct}$ method. A non-template control ($H_2O$) was used to confirm appropriate primer function.

**Primer Design**

Complementary DNA sequences for genes of interest were obtained from the NCBI website and primers were designed using the Primer3 website (https://primer3.ut.ee/) to the following criteria: $T_m$ 62.0°C, 18-23 bp in length, 40-60% GC content, 80-180 bp product size, no predicted self-complementarity or hairpins and one primer spanning an exon junction. Specificity of prospective primers to the gene of interest was confirmed using the Primer-BLAST online tool. Primer suitability was confirmed by preliminary tests involving a non-template control ($H_2O$) to check for specific amplification of the sequence of interest and the primer melt curves were inspected for anomalies such as primer dimers.

**Table S5.** Primer list Human (Homo sapiens)

| Primer | Sequence |
|---|---|
| ANG F1 | GCCTGGGCGTTTTGTTGTTG |
| ANG F2 | TCCTGGGAGCCTGTGTTGGA |

| | |
|---|---|
| ANG R1 | TGGGTCAGGAAGTGTGTGTACC |
| ANG R2 | GTATCTGTCATCCCGGCCCT |
| ANGPTL4 L | AGCCTGCCCGAAGAAAGAGG |
| ANGPTL4 R | GGCCGGTTGAAGTCCACTGA |
| APLN F | ACGCCCTCTTCCCTCTTGATG |
| APLN R | CAGCCCCACAACAGGACAGT |
| ARF1 F | TTCGCCAACAAGCAGGACCT |
| ARF1 R | CGCGTTCACTTCTGGTTCCG |
| ARF3 F | AGCAGGAACAGGATCAAGTCT |
| ARF3 R | CGCATCTCCTTCTTCCCAATC |
| ARF4 F | CTAGGGCTTCAGTCTCTTCGT |
| ARF4 R | ACAGCCAGTCAAGTCCTTCA |
| ARF5 F | CCACCATCCCAACCATAGGCT |
| ARF5 R | CGCTCCCGGTCATTACTGTC |
| ARG2 F1 | CCGTTCTCACAAGGGCAGAAAAG |
| ARG2 R1 | AGATCATCTTTGGGGACTGGAGT |
| CCND1 F | TGCCAACCTCCTCAACGACC |
| CCND1 R | TCCTCGCAGACCTCCAGCAT |
| CDH5 F | AGGTATGAGATCGTGGTGGAAGC |
| CDH5 R | TGTGTACTTGGTCTGGGTGAAGA |
| CDKN1A F | GTACCCTTGTGCCTCGCTCA |
| CDKN1A R | GCGTTTGGAGTGGTAGAAATCTG |
| CLDN3 F1 | CGCGAGAAGAAGTACACGGC |
| CLDN3 F2 | GGCCAACACCATTATCCGGG |
| CLDN3 F3 | CGCGAGAAGAAGTACACGGC |
| CLDN3 R1 | TAGTCCTTGCGGTCGTAGCC |
| CLDN3 R2 | GCCGTGTACTTCTTCTCGCG |
| CLDN3 R3 | TAGTCCTTGCGGTCGTAGCC |
| CXCL12 F | GTGCCCTTCAGATTGTAGCCC |
| CXCL12 R | CTCCTGAATCCACTTTAGCTTCG |
| CXCR4 F | ACGCCACCAACAGTCAGAGG |
| CXCR4 R | TTGGGGTAGAAGCGGTCACA |
| DDIT4 F | GTTCGCACACCCATTCAAGC |
| DDIT4 R | AGGCATGGTGAGGACAGACG |
| DUSP6 F | TGCTGGACTTCGAGAGGACG |
| DUSP6 R | AGATTGCAGAGAGTCCACCTGG |
| EDN1 F | ACAGACCGTGAAAATAGATGCCA |
| EDN1 R | TCTCCATAATGTCTTCAGCCCTG |
| EGLN3 F1 | TGCCCTGGAGTACATCGTGC |
| EGLN3 F2 | GCCCTCTTACGCAACCAGATATG |
| EGLN3 R1 | ATGCAGCGACCATCACCGTT |
| EGLN3 R2 | AGGGCAGATTCAGTTTTCCTAGT |

| | |
|---|---|
| ENO2 F | TGGAGAACAGTGAAGCCTTGGA |
| ENO2 R | TGGTCCCCAGTGATGTATCGG |
| ERRFI1 F | GTGATGAAGACAGGCCTCCCA |
| ERRFI1 R | AGGCTTTTGGGACTCGGTGT |
| F11R F | TCACAGTGCCTCCATCCAAGC |
| F11R R | CGGGTGCTTTTGGGATTCGT |
| FRAS1 F | GCACATCAGTTCATTGCCATCCA |
| FRAS1 R | GCACACACCATCCATTCTGCA |
| ICAM1 F | ACCATCTACAGCTTTCCGGC |
| ICAM1 R | CCATTCAGCGTCACCTTGGC |
| ID1 F | CAGGTAAACGTGCTGCTCTAC |
| ID1 R | AGAATCTCCACCTTGCTCACCT |
| IL6 F | TGAGAGTAGTGAGGAACAAGCCA |
| IL6 R | TTGTGGTTGGGTCAGGGGTG |
| ING4 F | GCTGAAATTGACAAGTTGGCCA |
| ING4 R | GTGTTTGTCCACCATCTCATAGG |
| LDHA F1 | AAGATTACAGTTGTTGGGGTTGG |
| LDHA F2 | ACAGTTGTTGGGGTTGGTGC |
| LDHA F3 | AGCAAGAGGGAGAAAGCCGT |
| LDHA R1 | AGTTCATCTGCCAAGTCCTTCAT |
| LDHA R2 | AGAGCAAGTTCATCTGCCAAGT |
| LDHA R3 | ACCACTTATCTTCCAAGCCACGT |
| MAP1LC3C F | AATCCCGGTGGTAGTGGAGC |
| MAP1LC3C R | ACCAGGCTCTTGTTGTTCACC |
| MTAP F | ACCGCCGTGAAGATTGGAAT |
| MTAP R | GTGCTGCCTTCCATGCCTTG |
| NOP56 F | TCCAGAGGAGTGTGAGGAGATGA |
| NOP56 R | TGCCAGCGGTCTCTTCAAGA |
| NOX4 F | AGGATCACAGAAGGTTCCAAGCA |
| NOX4 R | TGAGAAGTTGAGGGCATTCACCA |
| P27 F | ACCTGCAACCGACGATTCTTCT |
| P27 R | TGCTCCACAGAACCGGCATT |
| PGK1 F | AGCGGGTCGTTATGAGAGTCG |
| PGK1 R | TGGGACAGCAGCCTTAATCCT |
| PINK1 F | GCCTACATTGCCCCAGAACC |
| PINK1 R | GAGGAACCTGCCGAGATGTT |
| PNRC1 F | CGGAGAAAATTGCCCTTCCCC |
| PNRC1 R | GAGGTGATCTTGGTTAGTGGCA |
| PTGIS F | ATATGGGTCCCGCTGCCTTC |
| PTGIS R | TGTGGGAGAGTGGTCGTCTG |
| RAB33A F1 | CTGCCTGACCTTCCGCTTCT |
| RAB33A F2 | AGGGCGAGAAGATCAAGGTTCA |

| | |
|---|---|
| RAB33A R1 | TCCCACACCTGAACCTTGATCT |
| RAB33A R2 | CGTTGCGGTAGTAATGCTCGAC |
| RBM14 F | CCGCCGTTTATCAGAGTCGC |
| RBM14 R | CGTAATCGGAATGGGCATCGG |
| RPAP3 F | CCTGCAACCTCAAGCCAGTT |
| RPAP3 R | GCAGGAATTGGAGGAAGAACAGT |
| RRP9 F | AAGAAGCGACCACTTGCCCT |
| RRP9 R | TCTGTGTTGAGGAGGGCTGC |
| SELE F | TTCATGTTGCAGGGACCAGC |
| SELE R | ACTGGAAAGCTTCACAAACTGGG |
| SERPINE1 F1 | GGTGCTGGTGAATGCCCTCT |
| SERPINE1 F2 | CCACAAATCAGACGGCAGCA |
| SERPINE1 R1 | CGGGCGTGGTGAACTCAGTAT |
| SERPINE1 R2 | GGGCGTGGTGAACTCAGTATAGT |
| SLC2A1 F | GGGCCAAGAGTGTGCTAAAGA |
| SLC2A1 R | GGTGACCTTCTTCTCCCGCA |
| SLC39A12 F | TCCCTCAGGTTCTTGGTTTACAT |
| SLC39A12 R | TGCCTCCAATTAATCCCATCAGT |
| SOX17 F | TGAATCTCCCCGACAGCCAC |
| SOX17 R | TGTCACACGTCAGGATAGTTGCA |
| SPARC F | TGGATCTTCTTTCTCCTTTGCCT |
| SPARC R | CCCACAGATACCTCAGTCACCT |
| SQOR F | TGAGCCCAGTGAGAGACATTTC |
| SQOR R | TGGGTTCAACTCAGTCACTCTAG |
| TMEM201 F | GCACAGAACTTCTCCTCCGC |
| TMEM201 R | AAGTGTCCGCTGGCCCCATA |
| UPB1 F1 | TCATCTTCAACCCCTCGGCC |
| UPB1 F2 | TGGAAAGAAAGCTCACCAGGACT |
| UPB1 F3 | CGGGAGATGGAAAGAAAGCTCAC |
| UPB1 F4 | TGGAAAGAAAGCTCACCAGGACT |
| UPB1 R1 | AGTCCTGGTGAGCTTTCTTTCCA |
| UPB1 R2 | ACCTGCCCGTCATCTTGAAGT |
| UPB1 R3 | TCCAGACATCATTCACCTGCTG |
| UPB1 R4 | ACCTGCCCGTCATCTTGAAGT |
| VEGFA F | TGCCATCCAATCGAGACCCT |
| VEGFA R | CACTCCAGGCCCTCGTCATT |
| VEGFC F | GCCAATCACACTTCCTGCCG |
| VEGFC R | AGGTCTTGTTCGCTGCCTGA |

**Table S6.** Primer list mouse (Mus musculus)

| Primer | Sequence |
|---|---|
| Acvrl1 F | GTGTGTGGGAAAGGGCCGAT |
| Acvrl1 R | CCGGAACCAGGACTGCTCAT |
| Aldoc F | AATGGTGTCCCCTTCGTCCG |
| Aldoc R | CAAGAGCCCATCCAGCCCTT |
| Angptl4 F | GTTTGCAGACTCAGCTCAAGG |
| Angptl4 R | TCCATTGTCTAGGTGCGTGGG |
| Apln F | AGCCCAGAACTTCGAGGACT |
| Apln R | AGCCCTTCAATCCTGCTTTAGA |
| Aplnr F | GGTGCTCTGGACCGTGTTTC |
| Aplnr R | GGCCAGTCAAACTCCCGGTA |
| Arf1 F | TGACAGAGAGCGTGTGAACG |
| Arf1 R | CGCATTCATGGCATTGGGGA |
| Arf2 F | AGCTGGCAAAACGACGATCT |
| Arf2 R | GGCCACCAACATCCCAGACT |
| Arf3 F | TCTACAAGCTGAAACTCGGGGA |
| Arf3 R | CGAATCTTGTCCTGGCCACC |
| Arf4 F | TCTCTCCGAAACAGAACATGGTA |
| Arf4 R | ACAGCCAATCCAGTCCCTCA |
| Arf5 F | GCGGCACTACTTCCAGAACAC |
| Arf5 R | TCTGGAGTTCATCAGCAGACTC |
| Arf6 F | CGGGAACAAGGAAATGCGGA |
| Arf6 R | CCGATTGGCCCAGCTTCAAC |
| Arg1 F | TCCCTAATGACAGCTCCTTTCAA |
| Arg1 R | TGCTTCCAACTGCCAGACTGT |
| Arg2 F | TCCTCCACGGGCAAATTCCT |
| Arg2 R | AGCTTCTTCTGTCCCCGAGA |
| B2m F | GCCTGTATGCTATCCAGAAAACC |
| B2m R | TCAATGTGAGGCGGGTGGAA |
| Bmpr2 F | GTGATCCCCAAGAGTGCCACT |
| Bmpr2 R | AGGTGGACTGAGTGGTGTTGT |
| Bnip3l F | GGGACCACAGCTCTCAGTCAGA |
| Bnip3l R | GGGTGGGATGTTTTCGGGTCTA |
| Ca9 F | TTCCCTGCTGAGATCCACGT |
| Ca9 R | AGCCTTCCTCCGAGATTTCTTCC |
| Ccnd1 F2 | CGCGTACCCTGACACCAATC |
| Ccnd1 R2 | TGGCCACGATTTTCCGCATG |
| Cldn3 F | GCGGCTCTGCTCACCTTAGT |
| Cldn3 R | CACGTACAACCCAGCTCCCA |
| Cxcl12 F | GCCAACGTCAAGCATCTGAAAAT |

| | |
|---|---|
| Cxcl12 R | CGGGTCAATGCACACTTGTCTG |
| Cxcr4 F2 | TCTGTGACCGCCTTTACCCC |
| Cxcr4 R2 | TTCTGGTGGCCCTTGGAGTG |
| Dusp6 F | CTGTGCCAAGGACTCGACCA |
| Dusp6 R | ACTCGCCCGCATTCTCAAAC |
| Edn F | GCAGGAAAAGAACTCAGGGCC |
| Edn R | GGCCTCCAACCTTCGTAGTT |
| Egln3 F | GGGACGCCAAGTTACACGGA |
| Egln3 R | AGGGCTGGACTTCATGTGGA |
| Eln F | GCCAAATACGGAGCCAGAGG |
| Eln R | CCAACACCATAGCCAGGAAAGC |
| Eng F | CTCTACCTCAGCCCGCACTT |
| Eng R | AGACACGCTCACCTGTACGA |
| Eno2 F | CCAAAGGTCTTTTCCGGGCTG |
| Eno2 R | TCCTGCTGTTGATGTGGTCCA |
| Errfi1 F | CGGGACAGCGTGAAGAGGAT |
| Errfi1 R | CCGAGAACCCCGATCACACA |
| Fras1 F | TGGGTGTCCTCAAAGCGTGG |
| Fras1 R | TGATACAGACAAGCACCTTCACA |
| Grem1 F | TGAAGCAGACCATCCACGAGG |
| Grem1 R | GGTGAACTTCTTGGGCTTGCA |
| Icam1 F | AGCCTCCGGACTTTCGATCTT |
| Icam1 R | TTTGTGCTCTCCTGGGTCGG |
| Id1 F | GGCGAGGTGGTACTTGGTCT |
| Id1 R | GCTCACTTTGCGGTTCTGGG |
| Il6 F | ACTGGGGATGTCTGTAGCTCA |
| Il6 R | GCAACTGGATGGAAGTCTCTTGC |
| Ing4 F | GGCCCTTCTCAGACAGATCCA |
| Ing4 R | CGCCGAATGTGTTTGTCTACCAT |
| Ldha F | ATGAAGGACTTGGCGGATGAG |
| Ldha R | GCTTGGAGTTCGCAGTTACACA |
| Mtap F | GCCCCAAAACAAGAGAGGTCCT |
| Mtap R | GCGCCCCAAGTACGGAATAT |
| Nop56 F | TGGGGTACAACTGCCAGACTG |
| Nop56 R | TGTTGTCCACTCGGTTCACA |
| Nox4 F | CACCTCTGCCTGCTCATTTGG |
| Nox4 R | TCTGGCCCTTGGTTGTACAG |
| Pdk1 F | ACCAGCACTCCTTATTGTTCGGT |
| Pdk1 R | CTCCACCACGTCGCAGTTTG |
| Pgk1 F | GGTCGTGATGAGGGTGGACT |
| Pgk1 R | AACGGACTTGGCTCCATTGT |
| Prkg1 F | ACGTCATGGAAGATGGGAAGGT |

| | |
|---|---|
| Prkg1 R | ATAGCCAGCTCCCCGAACAC |
| Ptgis F | CGGTGACATATTTACTGTGCTGG |
| Ptgis R | TACGCAGCTCCCACACCACT |
| Rab33a F | ATCAAGGTTCAGGTGTGGGACA |
| Rab33a R | GGTGAAGGAGGTCATCTTGGTGA |
| Rrp9 F | GCGAAAGGTAGACTCTGCTGC |
| Rrp9 R | GCTCTCGCTCTCAGAATCGC |
| Sele F | CCCAACTGTGAGCAAGCTGTG |
| Sele R | ATAGCTGAAGGGGCCGAACG |
| Serpine1 F | TCTCTCCCTATGGCGTGTCC |
| Serpine1 R | TGTGCCCTTCTCATTGACTTTGA |
| Slc2a1 F | CCCCAGAAGGTTATTGAGGAGT |
| Slc2a1 R | GGCCACGGAGAGAGACCAAA |
| Slc39a12 F | TCTGAACATGCTCACGACCAGA |
| Slc39a12 R | GCAGAGAAGCAAGCCTGATCC |
| Vegfa F | AGCAGAAGTCCCATGAAGTGATC |
| Vegfa R | CAGGACGGCTTGAAGATGTACT |
| Ywhaz F2 | TCGCAACCAGAAAGCAAAGTCT |
| Ywhaz R2 | TGACTGGTCCACAATTCCTTTCT |

**Late outgrowth Endothelial Colony Forming Cells (ECFCs)**

Endothelial colony forming cells were derived from peripheral blood samples and characterised, as previously described (1, 7, 10). Venous blood samples were obtained with local ethics committee approval and informed written consent (REC Ref 17/LO/0563) from healthy volunteers (n=5) and PAH patients with rare pathogenic *BMPR2* variants (n=5). ECFCs were cultured on 0.2% gelatin (Sigma Aldrich, G1393) with the same endothelial medium as used for HPAECs, but with 20% HyClone FBS (Cytiva, SH30070.03) instead of the supplied FBS. All ECFCs were used between passages 3-6. Demographic and clinical features of healthy subjects and PAH patients are shown in **Table S7** below.

**Table S7.** Demographic and clinical features of healthy subjects and PAH patients

| Diagnosis | ID | Sex | Age | BMPR2 Variant |
|---|---|---|---|---|
| Healthy | C1 | Female | 57 | - |
| | C7 | Female | 23 | - |
| | C8 | Male | 22 | - |
| | C11 | Female | 29 | - |
| | C12 | Female | 36 | - |
| PAH | BP1 | Female | 65 | Frameshift |
| | BP2 | Male | 66 | Stop |
| | BP3 | Female | 37 | Stop |
| | BP4 | Male | 49 | Splice |
| | BP6 | Female | 43 | Missense |

**Study approval**

Venous blood samples were obtained with local ethics committee approval and informed written consent from healthy volunteers and idiopathic PAH patients. Participants were identified by number. Animal studies were conducted in accordance with UK Home Office Animals (Scientific Procedures) Act 1986.

**Apoptosis Assay**

Apoptosis was measured using Cell MeterTM Caspase 3/7 Activity Apoptosis Assay Kit (AAT Bioquest, ABD-22796). Fluorescence intensity was analysed in GlomaxTM luminometer at Ex/Em = 490/525 nm.

**Permeability Assay**

Changes in endothelial barrier function in HPAECs grown in Transwell-Clear filters (3-µm pore size, 12-mm diameter; Costar Corning, High Wycombe, UK) were assessed by measuring the passage of fluorescent dextran (FITC-dextran, MW 40 kDa, Sigma) through the endothelial cell layer, as previously described(7). Cells were grown to confluency prior to adenoviral infection. 2 hr post-infection the cells were left untreated or were treated with CTC, as required and placed under normoxic or hypoxic conditions and endothelial permeability was examined 24 hr later.

**Proliferation Assay**

HPAEC proliferation was studied in CyQUANT™ Cell Proliferation Assay (Thermofisher, # C7026), according to the manufacturer's protocol (https://www.thermofisher.com/document-connect/document-connect.html?url=https://assets.thermofisher.com/TFS-Assets%2FLSG%2Fmanuals%2Fmp35006.pdf)

**Receptor Internalisation Assay**

To assess receptor internalisation, HPAECs were treating with ARF6 mutant adenoviruses for 24 hours as described above. Next, cells were placed on ice and washed with ice-cold PBS. The cell surface was then labelled with 380µg/ml EZ-Link Sulfo-NHS-SS-Biotin (Thermo Fisher, A39258) in ice-cold PBS for 30 min at 4°C, and unbound biotin was removed with two washes of ice-cold PBS. Media was then added and cells were replaced at 37°C for 30 minutes, to allow internalisation to occur. Cells were then placed immediately on ice and washed with cold PBS. Biotin remaining at the cell surface after internalization was removed with 60mM MES-Na (Merck, M1511) in reduction buffer (50mM Tris pH 8.6, 100 mM NaCl) for 30 minutes on a rocker at 4°C followed by quenching with 100 mM iodoacetamide (Sigma Aldrich, I1149) for 10 minutes, leaving only internalised biotin inside the cells. This step was omitted from one well to allow for quantification of total biotin load, so that the relative amounts of biotinylated internalised proteins were normalized to the biotin signal of all labelled proteins. Internalised VE-cadherin was fluorescently labelled using an Alexa-488 conjugated antibody (1:250, Thermo Fisher, 53-1449-42) in 30mM HEPES for 1 hour on ice. This was then quantified on a plate reader at 480/525nm Ex/Em. This protocol was adapted from (11).

**SUPPLEMNTAL TABLES (RESULTS)**

**Table S8. Differentially expressed proteins by proteomics.** WT: wildtype ARF6, CA: constitutive active ARF6, DN: dominant negative ARF6.

| ARF6 Mutant | Protein | Fold-Change | P value | q value |
|---|---|---|---|---|
| WT | ARF6 | 3.63 | 0.00055 | 0.03567 |
| | NRP2 | 1.53 | 0.00056 | 0.03567 |
| | EFNB1 | 1.26 | 0.00001 | 0.00603 |
| CA | NRP2 | 1.57 | 0.00042 | 0.02175 |
| | PROCR | 1.36 | 0.00051 | 0.02350 |
| | SHE | 1.24 | 0.00096 | 0.03225 |
| | EPHB2 | 1.24 | 0.00017 | 0.01331 |
| | CGNL1 | 1.23 | 0.00048 | 0.02249 |
| | EPHB4 | 1.23 | 0.00212 | 0.04886 |
| | CDH11 | 1.23 | 0.00031 | 0.01878 |
| | CDK1 | 0.80 | 0.00014 | 0.01331 |
| | MRTO4 | 0.78 | 0.00073 | 0.02692 |
| | DDX21 | 0.78 | 0.00190 | 0.04582 |
| | CCN2 | 0.75 | 0.00218 | 0.04893 |
| | DNMT1 | 0.73 | 0.00227 | 0.04999 |
| | GNL2 | 0.70 | 0.00115 | 0.03445 |
| | H1-2 | 0.66 | 0.00178 | 0.04575 |
| FC | ARF6 | 3.46 | <0.000001 | 0.00012 |
| | PROCR | 1.57 | 0.00012 | 0.01261 |
| | EFNB1 | 1.56 | 0.00046 | 0.03138 |
| | SLC2A3 | 1.41 | 0.00001 | 0.00227 |
| | NID1 | 1.38 | 0.00111 | 0.04699 |
| | TSC2 | 1.29 | 0.00059 | 0.03577 |

| | | | | |
|---|---|---|---|---|
| | CD276 | 1.26 | 0.00000 | 0.00157 |
| | LPCAT1 | 1.21 | 0.00014 | 0.01404 |
| | MRTFB | 0.80 | <0.000001 | 0.00058 |
| | TBC1D15 | 0.79 | 0.00015 | 0.01404 |
| | FYN | 0.79 | 0.00072 | 0.03974 |
| | POLR2A | 0.76 | 0.00011 | 0.01261 |
| | RRP15 | 0.76 | 0.00036 | 0.02566 |
| | CRIM1 | 0.73 | 0.00101 | 0.04686 |
| | DHFR | 0.67 | 0.00108 | 0.04699 |
| | RAD50 | 0.67 | 0.00008 | 0.00992 |
| | MRE11 | 0.65 | 0.00002 | 0.00435 |
| | CCN2 | 0.64 | 0.00001 | 0.00227 |
| | GJA1 | 0.64 | 0.00078 | 0.04177 |
| **DN** | ARF6 | 6.02 | 0.00030 | 0.01556 |
| | NRP2 | 1.80 | 0.00121 | 0.02713 |
| | FADS2 | 1.72 | 0.00087 | 0.02325 |
| | PROCR | 1.59 | 0.00003 | 0.00483 |
| | EFNB1 | 1.46 | 0.00040 | 0.01760 |
| | CDH11 | 1.45 | 0.00011 | 0.00899 |
| | DCBLD2 | 1.39 | 0.00086 | 0.02318 |
| | EPHB4 | 1.39 | 0.00154 | 0.03146 |
| | BCAR3 | 1.37 | 0.00078 | 0.02289 |
| | GPC1 | 1.34 | 0.00218 | 0.03987 |
| | SHE | 1.32 | 0.00285 | 0.04613 |
| | HYAL2 | 1.28 | 0.00003 | 0.00522 |
| | IDI1 | 1.26 | 0.00006 | 0.00627 |
| | EIF1AD | 1.25 | 0.00103 | 0.02481 |
| | ADAM15 | 1.24 | 0.00151 | 0.03132 |

| | | | | |
|---|---|---|---|---|
| | PEAR1 | 1.23 | 0.00059 | 0.02002 |
| | CD276 | 1.23 | 0.00091 | 0.02379 |
| | NADSYN1 | 1.21 | 0.00000 | 0.00093 |
| | MED20 | 1.21 | 0.00043 | 0.01821 |
| | STOM | 1.21 | 0.00209 | 0.03871 |
| | ALCAM | 1.20 | 0.00049 | 0.01873 |
| | SMC4 | 0.80 | 0.00291 | 0.04678 |
| | HK2 | 0.79 | 0.00229 | 0.04116 |
| | ERI1 | 0.79 | 0.00120 | 0.02713 |
| | FTO | 0.78 | 0.00269 | 0.04488 |
| | RRP1 | 0.75 | 0.00084 | 0.02318 |
| | TP53BP1 | 0.75 | 0.00016 | 0.01123 |
| | HAUS8 | 0.75 | 0.00162 | 0.03217 |
| | VRK1 | 0.74 | 0.00003 | 0.00522 |
| | MCM5 | 0.73 | 0.00278 | 0.04574 |
| | POLR2A | 0.72 | <0.000001 | 0.00036 |
| | GIMAP7 | 0.72 | 0.00070 | 0.02161 |
| | CRIM1 | 0.69 | 0.00192 | 0.03687 |
| | SLC43A3 | 0.69 | 0.00011 | 0.00908 |
| | CDK1 | 0.68 | 0.00013 | 0.00949 |
| | MRE11 | 0.67 | 0.00242 | 0.04284 |
| | NDC80 | 0.67 | 0.00012 | 0.00920 |
| | RAD50 | 0.66 | 0.00080 | 0.02299 |
| | PBK | 0.65 | 0.00146 | 0.03061 |
| | GJA1 | 0.64 | 0.00154 | 0.03146 |
| | SPC24 | 0.61 | 0.00297 | 0.04710 |
| | DHFR | 0.61 | 0.00037 | 0.01744 |
| | CCN2 | 0.59 | 0.00061 | 0.02025 |

**SUPPLEMNTAL FIGURES        Fellows et al.**

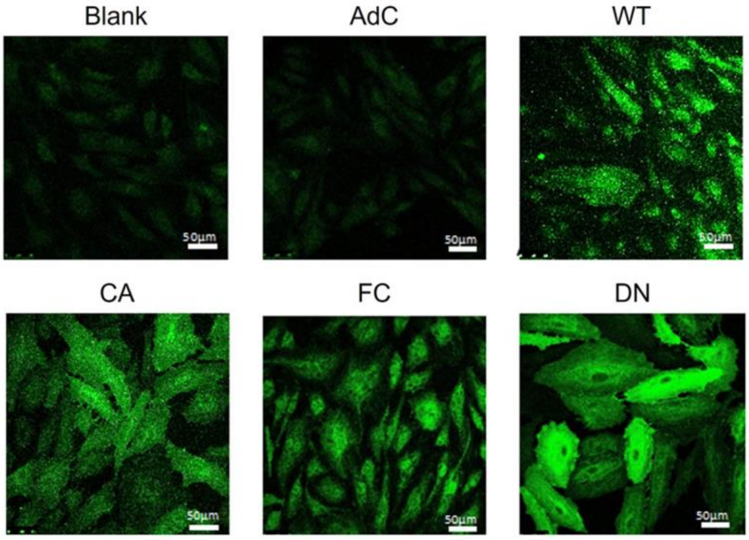

**Figure S1. Cellular distribution of overexpressed ARF6 mutant proteins.**

Fluorescence microscopy images of HPAECs 24 hours after adenoviral infection, with total ARF6 shown in green (immunofluorescence).  AdC - adenoviral control; WT - wildtype, CA – constitutively active, FC -fast cycling, DN – dominant negative Arf6 mutant. Bar=50 µm

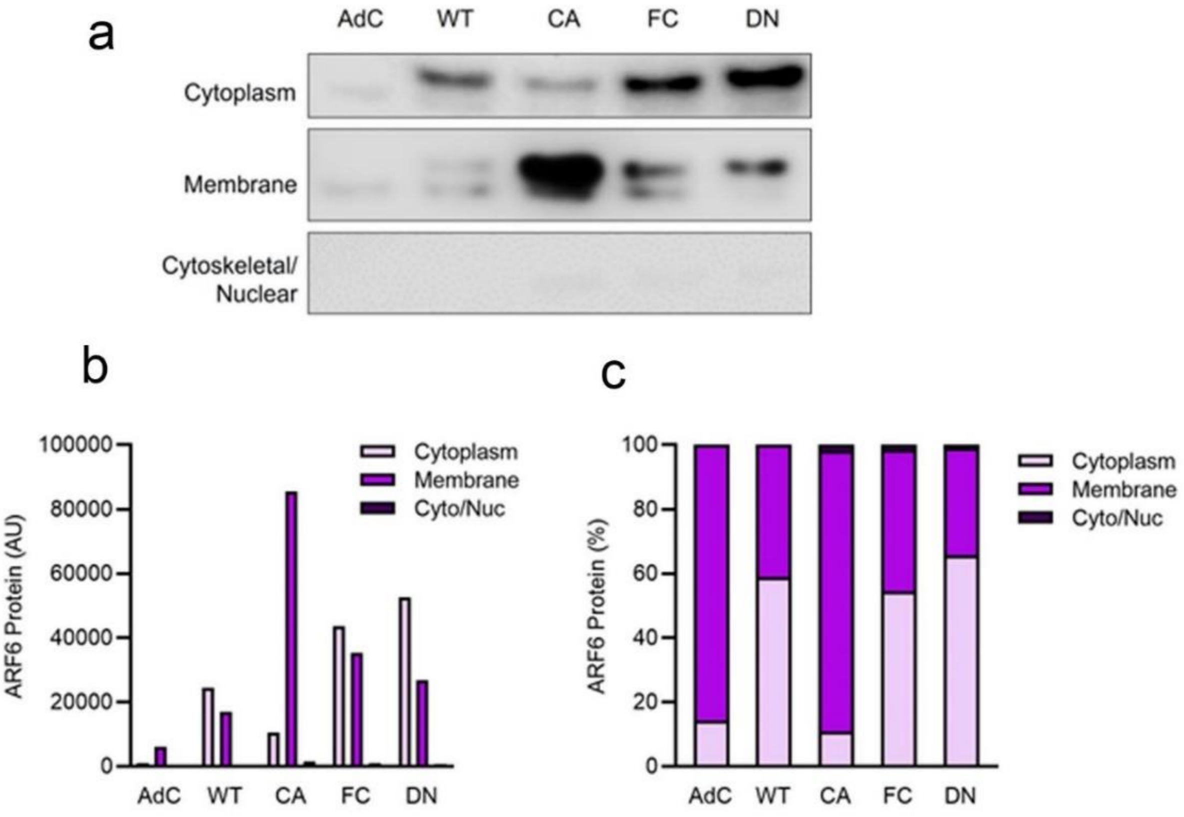

**Figure S2. Cellular localisation of overexpressed ARF6 mutant proteins.**
(a) Western blots showing ARF6 levels in cytoplasmic, membrane and cytoskeletal/nuclear fractions of HPAECs, 24 hours after adenoviral transduction. (b) Quantification of ARF6 protein levels in different fractions. (c) Distribution of ARF6 in different fractions expressed as a percentage of total ARF6.

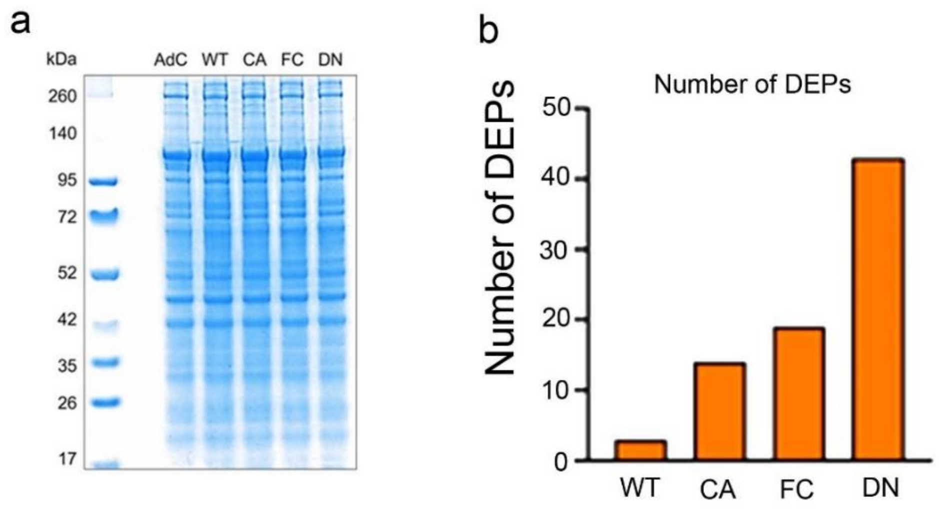

**Figure S3. Impact of ARF6 mutants on intracellular proteins.**

(a) Coomassie blue staining of a polyacrylamide gel following SDS-PAGE of HPAEC lysates 24 hours after adenoviral transduction with control (AdC) and WT, CA, FC and DN mutants of Arf6, as indicated. (b) Number of differentially expressed proteins (DEPS) in HPAECs overexpressing Arf6 mutants.

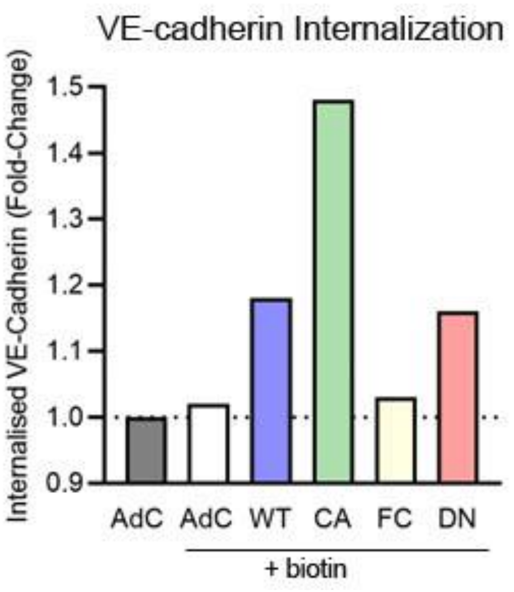

**Figure S4. Effect of ARF6 mutants on membrane internalisation of VE-cadherin.**

Internalisation assay showing levels of intracellular, biotin-labelled VE-cadherin measured by ELISA. 'AdC' indicates antibody control (without biotin) and +' indicates measurements with biotinylated antibody; fold-change of control.

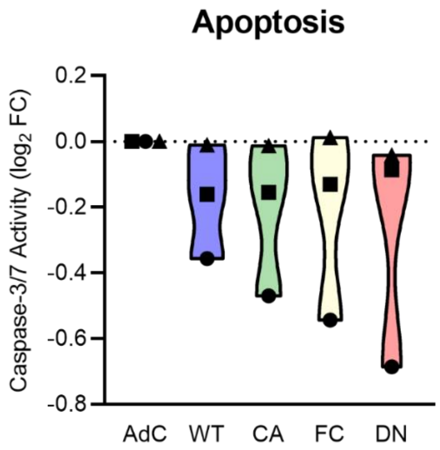

| Dunnett's multiple comparisons test | Adjusted P Value |
|---|---|
| AdC vs. WT | 0.789 |
| AdC vs. CA | 0.674 |
| AdC vs. FC | 0.648 |
| AdC vs. DN | 0.489 |

**Figure S5**. **Effect of ARF6 activity mutants on cell apoptosis.**
Caspase-3/7 assay 24 hours after adenoviral transduction of HPAECs with different Arf6 activity mutants, as indicated (n=3). One-way ANOVA with Dunnett's post-test.

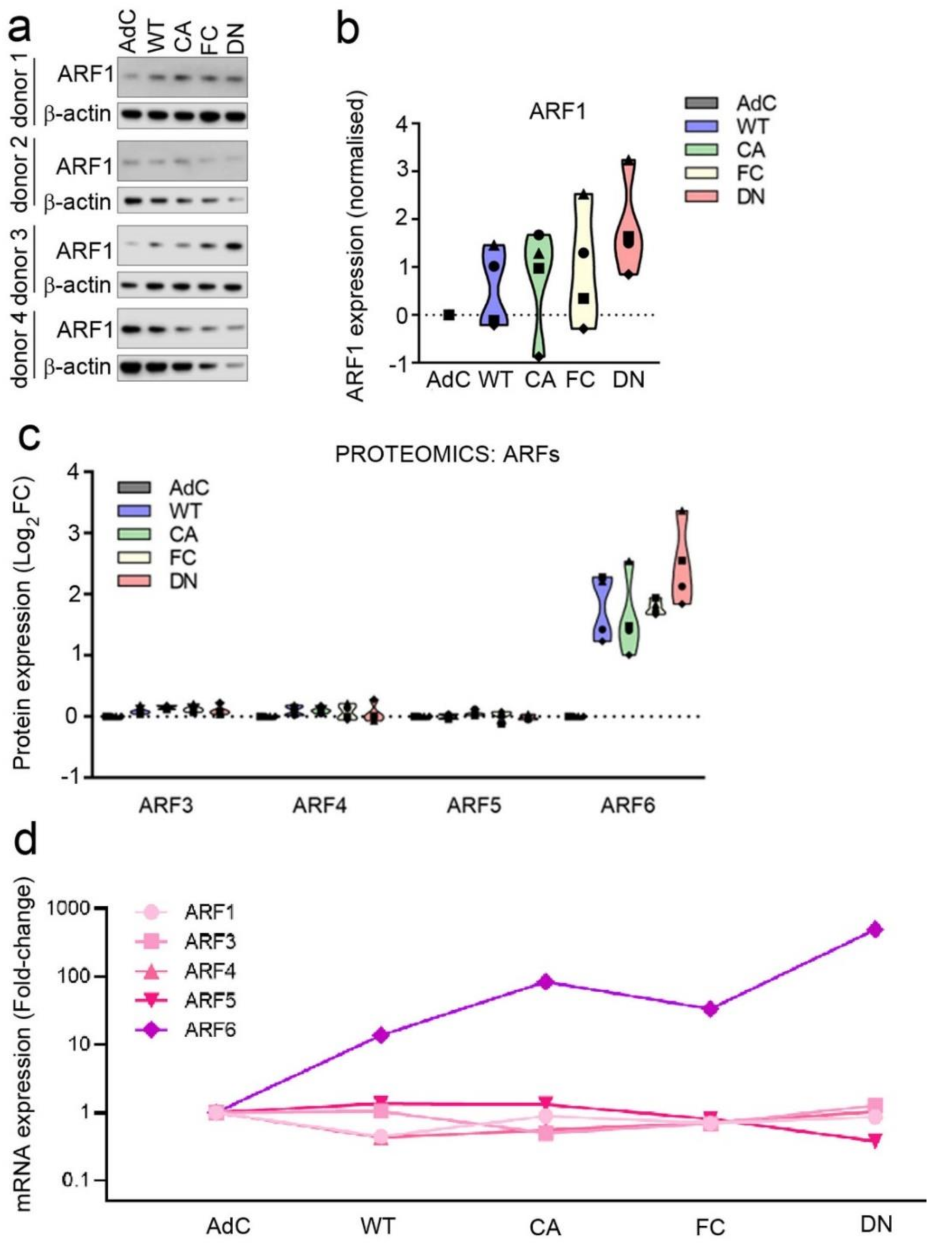

**Figure S6. ARFf6 activity mutants do not affect expression of other ARF proteins.**

(a) Western blots and (b) a corresponding graph show effect of Arf6 activity mutants on protein expression of ARF1 24 hour post-infection. (c) ARF6 protein levels from proteomic analysis and (d) mRNA levels measured by qPCR in HPAECs overexpressing ARF6 mutants for 24 hours and 16 hours, respectively. In a-c, n=4; in d, n=1.

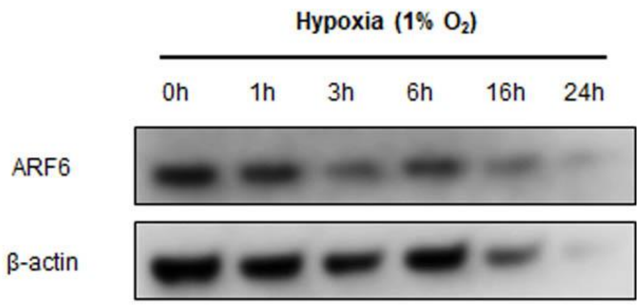

**Figure S7**. **Hypoxia does not affect Arf6 protein expression.**

HPAECs were subjected to hypoxia for 0-24h, as indicated and Arf6 expression was evaluated by western blotting.

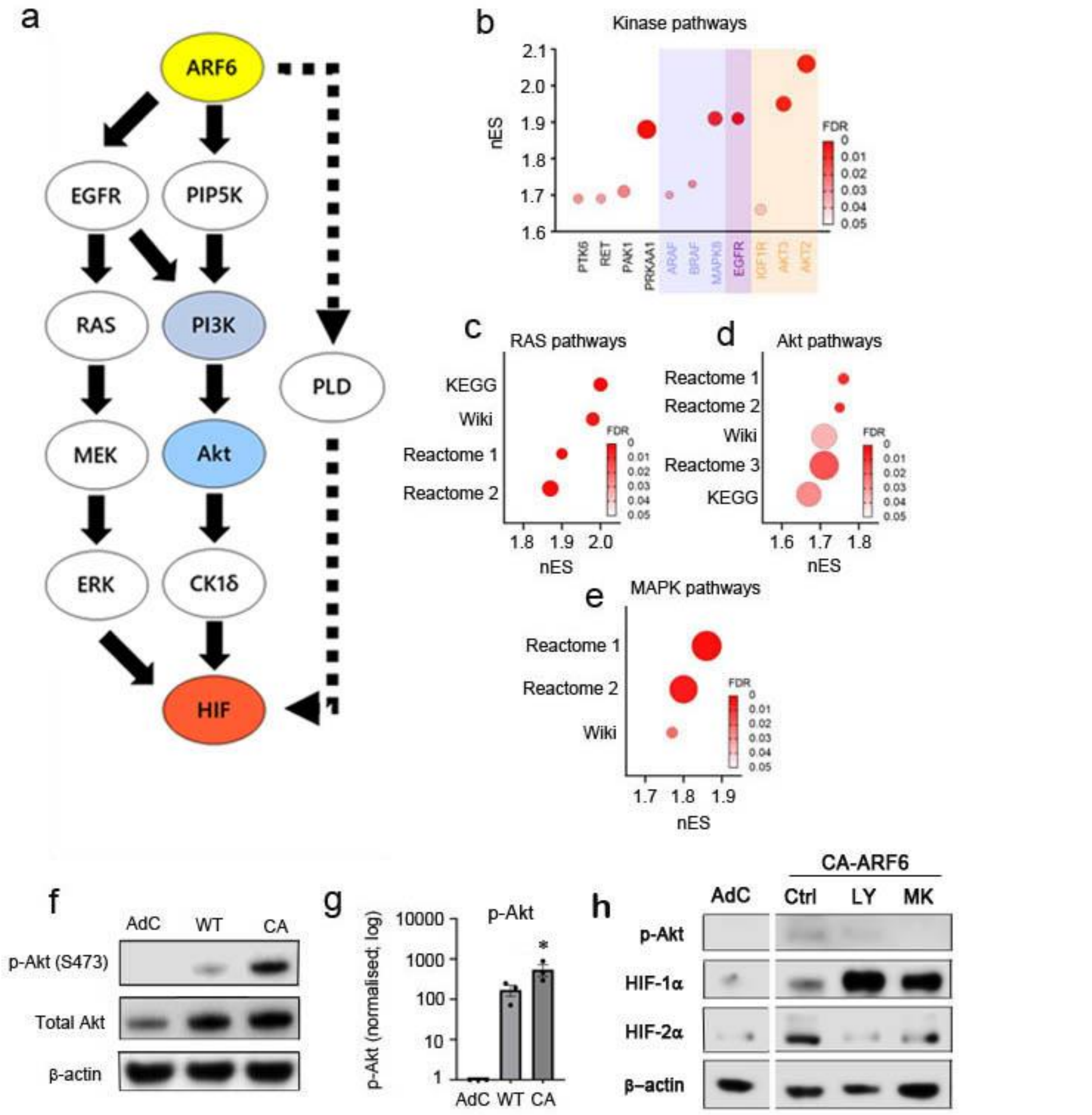

**Figure S8. Proposed mechanism of ARF6-mediated HIF activation.**
(a) Schematic of proposed ARF6-HIF signalling. For further detail, please see Discussion. (b) Bubble plot showing CA-ARF6-upregulated kinases, those from similar pathways are highlighted with the same colours. (c) Bubble plot showing enrichment of pathways referencing ‘RAS’ from different bioinformatics databases by CA-ARF6. (d) Bubble plot showing enrichment of pathways referencing ‘Akt’ from different bioinformatics databases by CA-ARF6. (e) Bubble plot showing enrichment of pathways referencing ‘MAPK’ from different bioinformatics databases by CA-ARF6. (f) Western blots and (g) a corresponding graph showing quantification of phosphorylated (active) and total Akt in HPAEC whole cell lysates following overexpression of ARF6 mutants. In the graph, p-Akt levels were normalised to total Akt expression; (n=4). (h) expression of p-Akt, HIF-1α and HIF-2α in control (AdC) and Ad-CA-ARF6-overexpressing HPAECs. The cells were left untreated (Ctrl) or were treated with Akt inhibitor (MK) and PI3K inhibitor (LY), as indicated. *P<0.05. one way ANOVA, comparison with AdC.

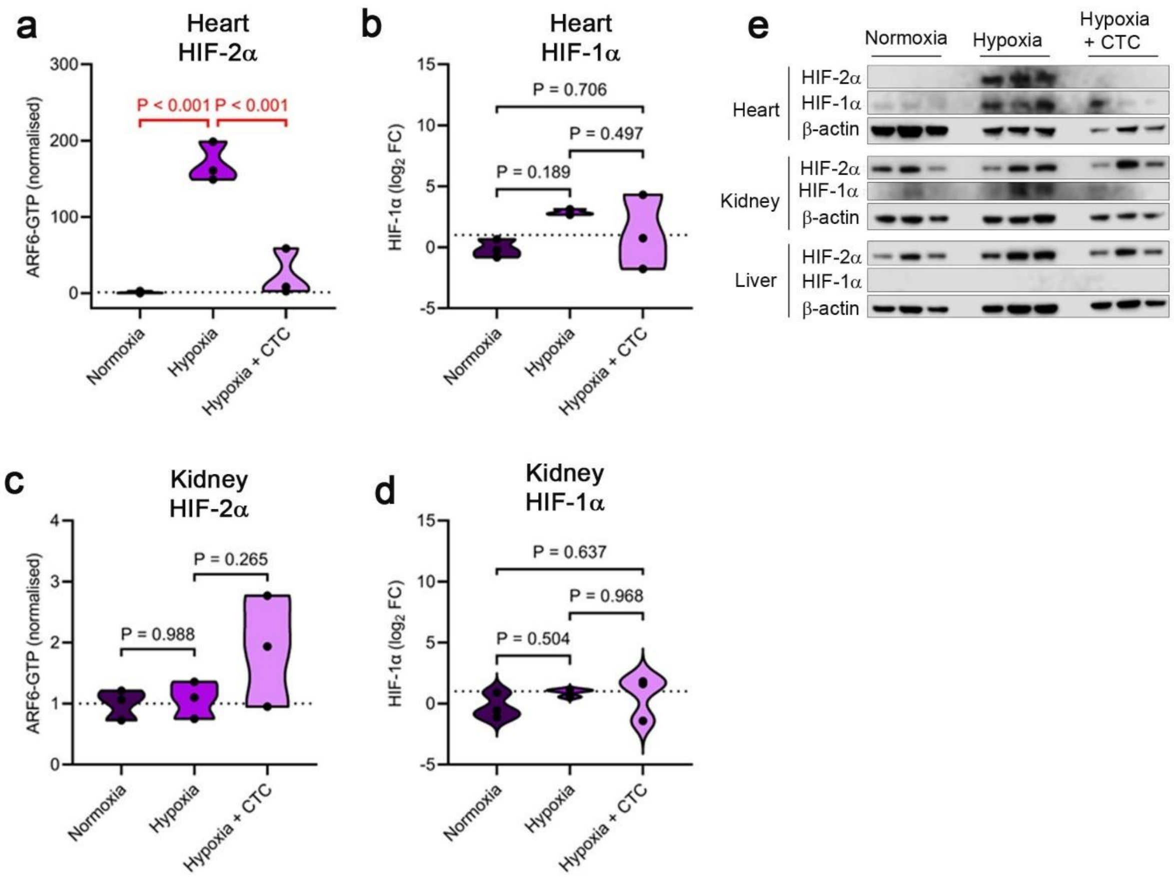

**Figure S9. Effects of CTC on hypoxia-induced HIF-1α and HIF-2α expression are tissue-specific.** Graphs in (a-d) show protein expression of HIF-1α and HIF-2α in heart and kidney tissues of normoxic mice, hypoxic mice (24h hypoxia) and hypoxic mice treated with CTC, as indicated. (e) corresponding representative western blots showing HIF levels in heart, kidney and liver. n=3. P values are, as indicated, one way ANOVA- with Tukey post-test.

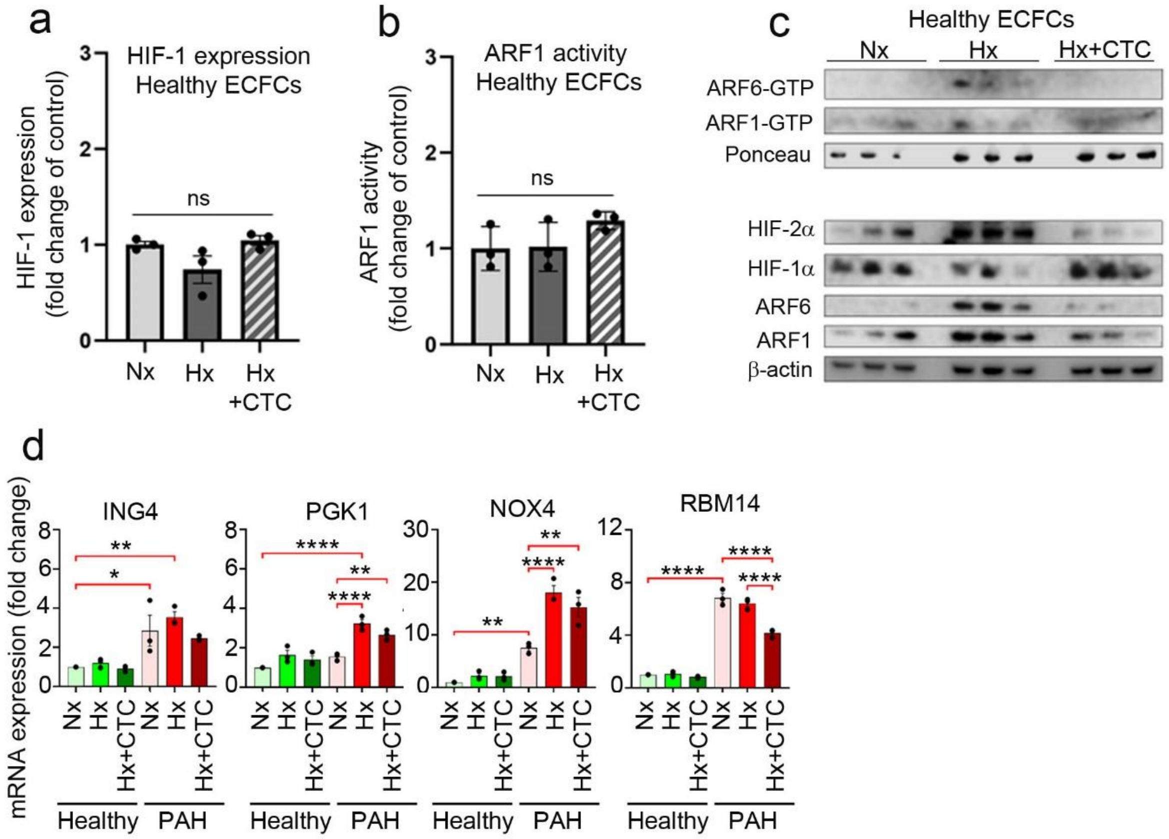

**Figure S10. Effect of chlortetracycline (CTC) on ARF6 and HIF activity in healthy and PAH ECFCs**.

Changes in (a) HIF-1 expression; (b) ARF1 activity in healthy ECFCs cultured under normoxic (Nx) or hypoxic (Hx) contridions and treated with CTC, as indicated. (c) corresponding westen blots and (d) expression of selected HIF-regulated genes in healthy and PAH ECFCs cultured under normoxic or hypoxic conditions, as indicated. In (a, b) bars are means ± SEM, ns-non significant; . In (d) *P<0.05, **P<0.01, ***P<0.001, comparisons, as indicated; one way ANOVA with Tukey post-test, n=3 (3 different donors).
